# Spatium: A Protein Language Foundation Model for Spatial Proteomics

**DOI:** 10.64898/2026.07.23.740264

**Authors:** Tiangang Wang, Sijia Wu, Liyu Huang, Jiajia Liu, Kexin Huang, Xiaobo Zhou

## Abstract

Spatial proteomics provides single-cell protein measurements under highly constrained and heterogeneous protein panels across datasets, resulting in limited and partially overlapping measurement spaces for cellular characterization. Existing analyses predominantly rely on statistical or task-specific modeling, while learning scalable representations of spatial protein data remain underexplored. This gap motivates the need for models that can learn stable representations of cellular identity from constrained protein measurements. Here we introduce Spatium, a protein language foundation model trained on over 51 million cells across multiple spatial proteomics platforms. Spatium learns intrinsic co-expression hierarchies that capture cell identity in a manner robust to panel composition and measurement scale. Spatium builds a generalizable representation of cell states grounded in biologically interpretable protein expression patterns. We demonstrate that Spatium learns biologically meaningful cell representations across multiple downstream tasks. Spatium recovers accurate cell identities with marker expression patterns consistent with known biology and reveals functionally distinct spatial microenvironments characterized by coherent marker enrichment signatures. It further enables reconstruction of missing protein measurements while preserving biologically meaningful expression patterns. Across these analyses, Spatium demonstrates stable and interpretable performance with lightweight task-specific adaptation, highlighting the robustness of the learned representations across diverse biological and experimental contexts.

## Introduction

Unlike spatial transcriptomics, which profiles gene expression across tissue sections[1], spatial proteomics enables the simultaneous measurement of dozens of protein markers at single-cell resolution, directly capturing cell identity and tissue architecture through their molecular phenotypes[2]. By preserving the spatial coordinates of each cell within intact tissue, these technologies provide a spatially resolved view of cellular organization, intercellular interactions, and microenvironmental structure[3]. Major platforms, including CODEX[4], IMC[5], CyCIF[6], and MIBI[7], have been widely applied to characterize tumor microenvironments, immune cell organization, and tissue structure, generating large-scale datasets that capture the molecular and spatial complexity of tissues across diverse disease contexts[8, 9]. The development of these platforms has resulted in the accumulation of large-scale spatial proteomics datasets, encompassing millions of spatially resolved single cells across diverse tissues and disease states[10–12]. The increasing scale and diversity of spatial proteomics datasets motivate the development of foundation models that can learn generalizable representations of cellular identity across tissues and disease contexts.

Foundation models have recently enabled powerful representation learning in single-cell genomics by capturing structured patterns of gene expression underlying cellular identity[13–15]. However, spatial proteomics differs fundamentally in its measurement regime: cellular identity is encoded through a limited set of protein markers whose combinatorial expression patterns and spatial context jointly define cellular states[8, 16]. This results in a representation setting where cellular identity must be inferred from a limited set of protein markers with strong combinatorial structure, rather than from high-dimensional continuous gene expression signals[17, 18]. However, most existing methods for spatial proteomics remain largely task-specific or statistical in nature[8, 19], lacking a unified framework for learning scalable representations of cellular identity from protein-level measurements.

Spatial proteomics exhibits a strong underlying biological structure, where cells of the same type are characterized by highly consistent combinations of protein markers, reflecting the fact that marker panels are designed for functional and phenotypic identification. Despite this inherent structure, existing computational approaches primarily operate in a task-driven manner. For example, methods such as MaxFuse[19], STELLAR[20], and scMODAL[21] focus on cross-modality alignment and label transfer, while marker-based approaches like CELESTA[22] rely on anchor matching for cell type annotation. In addition, clustering-based methods such as CytoCommunity[23] and manual neighborhood analysis typically depend on predefined markers or downstream labels rather than learning latent representations. Even recent foundation-style models such as KRONOS[24] operate on imaging-based representations and focus on morphological rather than protein-expression-driven modeling. Collectively, these approaches remain rooted in task-specific or heuristic designs, lacking a unified representation learning framework that can capture shared protein expression structure across cells and tissues in a common latent space. This limits their ability to model the underlying structure of cellular phenotypes in a consistent manner across spatial contexts. This highlights the need for a foundation model that learns generalizable representations of cellular identity from spatial protein data.

Here, we present Spatium, a protein language foundation model trained on large-scale spatial proteomics data spanning over 50 million cells across multiple platforms, including CODEX, IMC, CyCIF, and MIBI. Spatium learns representations of cellular identity through a self-supervised masked protein modeling objective in a BERT-style framework[25] applied to protein expression profiles, enabling the model to learn contextual dependencies among proteins and encode cellular states in a structured representation space. Importantly, the learned representations capture both discrete cell type organization and continuous phenotypic variation within cell populations, enabling fine-grained characterization of cellular subtypes beyond coarse annotations. Without requiring explicit full supervision, these representations can be effectively adapted to downstream tasks through lightweight task-specific heads, supporting accurate cell type annotation, spatial niche identification, and protein expression imputation. In addition, Spatium enables the discovery of functionally coherent spatial microenvironments characterized by consistent cellular composition and spatial organization, facilitating the identification of biologically meaningful tissue niches. The model is robust to variability in protein panels and measurement sparsity across datasets, allowing it to infer consistent cellular identities even under incomplete or partially overlapping marker sets. Furthermore, Spatium exhibits adaptability across experimental settings, including cross-platform and cross-cohort scenarios, demonstrating its potential as a general-purpose foundation model for spatial proteomics. Collectively, these capabilities establish Spatium as a unified representation learning framework for spatial protein data, capable of capturing shared biological structure across cells, tissues, and measurement contexts.

## Results

### Overview of Spatium

Spatial proteomics provides high-dimensional measurements of protein expression while preserving cellular localization within tissues. However, existing spatial proteomics datasets are generated using diverse imaging platforms, tissue contexts and marker panels, resulting in substantial heterogeneity that limits direct integration and large-scale representation learning[26]. To address this challenge, we developed Spatium, a foundation model designed to learn transferable protein-state representations from heterogeneous spatial proteomics datasets.

Spatium was pretrained on a large-scale compendium comprising 51 million cells from 70 human spatial proteomics studies, encompassing diverse tumor, immune and stromal compartments. As illustrated in Fig. 1a, spatial proteomics images were first converted into cell-level protein expression matrices through image segmentation and quantification of protein intensities[27]. Each cell was represented as a protein expression profile together with contextual metadata, including technology, tissue type and disease state. These cell-level representations were subsequently transformed into unified protein token sequences and processed by a Transformer-based encoder to learn generalizable representations of cellular protein states.

**Fig. 1.**
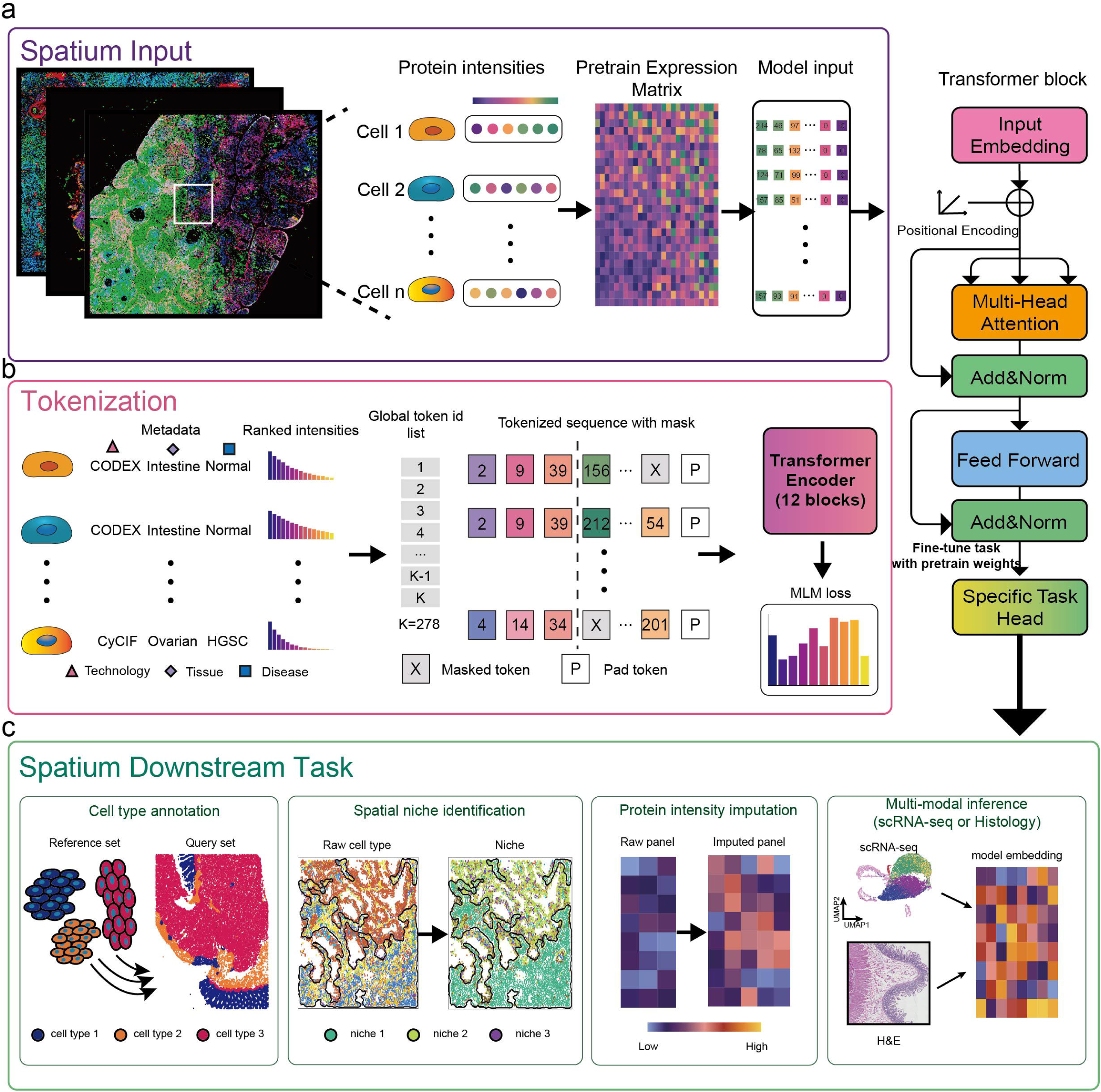
Spatium: a foundation model for spatial proteomics. (a). Overview of the Spatium pretraining framework. Multiplexed spatial proteomics images are first segmented to obtain single-cell protein intensities, which are assembled into cell-by-protein expression matrices. Each cell is subsequently transformed into a cell sequence and used as input to a Transformer encoder for self-supervised pretraining. (b). Rank-based protein tokenization and masked language modeling (MLM) pretraining. To enable joint learning across heterogeneous spatial proteomics datasets with different protein panels, protein names are first standardized into a unified protein vocabulary. Within each cell, proteins are ordered according to their normalized expression levels and converted into ranked protein tokens. Metadata describing imaging technology, tissue type and disease status are incorporated as side-information tokens together with a classification token 〈*cls*〉. Protein sequences are padded to a fixed length and randomly masked following a BERT-style masking strategy. The Transformer encoder is pretrained to recover masked protein identities from their surrounding ranked protein context using an MLM objective, thereby learning contextual representations of cellular protein programs that are independent of dataset-specific panel composition and absolute fluorescence intensities. (c). Overview of downstream applications supported by Spatium. The pretrained model provides transferable cellular representations that can be adapted through task-specific fine-tuning for diverse spatial proteomics analyses. Cell type annotation transfers cellular identity from annotated reference datasets to unlabeled spatial datasets. Spatial niche identification incorporates local neighborhood composition to discover higher-order tissue microenvironments. Protein intensity imputation reconstructs missing protein measurements from incomplete marker panels using contextualized protein representations. The learned embedding space further supports multimodal inference by integrating spatial proteomics with complementary modalities, including single-cell RNA sequencing and histopathology imaging.

A central challenge for spatial proteomics foundation models is the substantial heterogeneity among experimental panels[26]. Different studies frequently measure different protein subsets, use distinct naming conventions for equivalent markers, and exhibit dataset-specific intensity distributions. Furthermore, spatial proteomics panels are typically designed as targeted marker collections rather than comprehensive proteomic measurements, resulting in sparse and incomplete feature spaces.

To overcome these limitations, Spatium introduces a rank-based protein tokenization strategy (Fig. 1b). Protein identities across studies were first harmonized into a unified protein vocabulary. Rather than relying on absolute expression values, proteins within each cell were ordered according to their relative abundance, generating a ranked protein sequence that captures the internal organization of cellular protein states while reducing sensitivity to dataset-specific intensity variation[13, 14, 28].

For each cell, ranked proteins were mapped into a shared token vocabulary and combined with auxiliary metadata tokens describing experimental context. The resulting sequence consisted of metadata tokens, special tokens and ranked protein tokens, with missing positions padded to a fixed sequence length. During pretraining, Spatium was optimized using a masked language modeling objective, where randomly masked protein tokens were predicted from the surrounding protein context[14, 25]. This self-supervised learning strategy enables the model to capture combinatorial protein patterns that define cellular identity while remaining robust to incomplete measurements.

The representations learned by Spatium provide a transferable embedding space that captures cellular protein states and their spatial context, enabling flexible adaptation to diverse spatial biology tasks (Fig. 1c). By preserving both molecular identity and contextual information, these embeddings can be transferred across datasets and experimental settings, supporting consistent interpretation of previously unseen spatial proteomics data. For example, Spatium embeddings enable cell-state annotation by transferring cellular identity information from reference datasets, while incorporation of neighborhood information allows characterization of spatially organized tissue ecosystems beyond individual cell identities. The same representation space can further support reconstruction of missing protein measurements from incomplete panels and facilitate alignment with complementary modalities, including single-cell transcriptomics and histological imaging. Together, these applications highlight the ability of Spatium to serve as a general-purpose representation model for spatial proteomics, providing a unified framework for extracting transferable biological information from heterogeneous spatial datasets.

### Large-scale pretraining enables flexible adaptation of Spatium representations across spatial proteomics tasks

Foundation models rely on large-scale and diverse training corpora to learn transferable representations that capture shared biological principles across heterogeneous datasets[13–15, 28, 29]. To establish a generalizable representation framework for spatial proteomics, we assembled a comprehensive pretraining corpus consisting of 70 spatial proteomics studies, encompassing 51.8 million cells across four major imaging-based technologies, including CODEX, CyCIF, IMC and MIBI (Fig. 2b, c, d). These datasets span 18 tissue types and 18 disease contexts, covering diverse biological states ranging from normal tissues to multiple tumor and inflammatory conditions. To enable representation learning across studies with distinct experimental designs and marker panels, protein measurements were harmonized into a unified vocabulary of 239 protein markers.

**Fig. 2.**
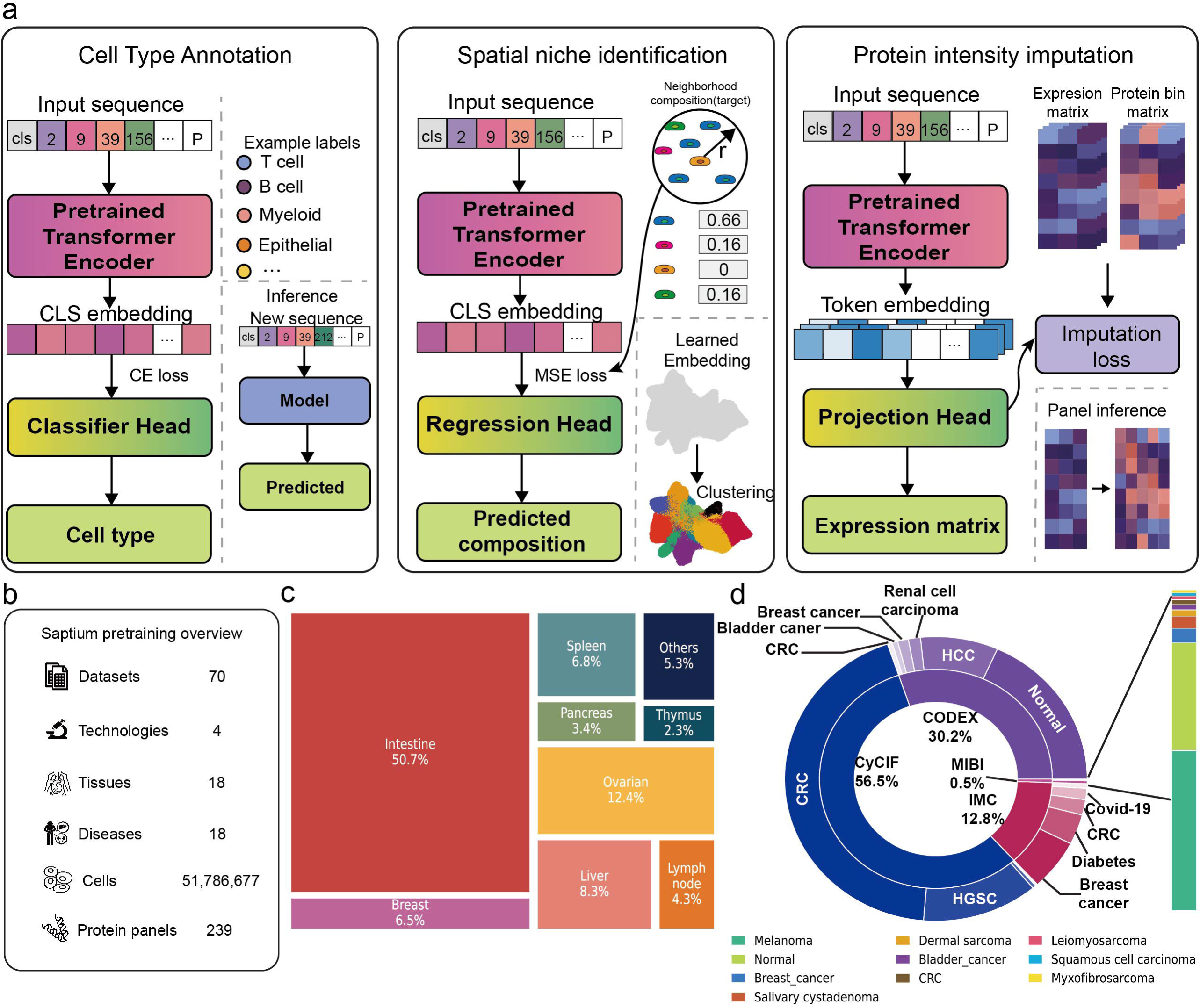
Fine-tuning strategies and pretraining data overview of Spatium. (a). Task-specific fine-tuning strategies built upon the pretrained Spatium encoder. The same pretrained backbone is adapted for cell type annotation, spatial niche identification and protein intensity imputation by attaching lightweight task-specific prediction heads while preserving the shared pretrained representation. (b). Summary of the spatial proteomics corpus used for Spatium pretraining, including the total number of datasets, single cells, imaging technologies, tissues, diseases and the unified protein vocabulary. (c). Distribution of pretraining cells across tissue types. The tree map summarizes the relative contribution of each tissue to the complete pretraining corpus. (d). Composition of the pretraining dataset across imaging technologies and disease types. The inner ring indicates the proportion of cells contributed by each spatial proteomics technology (CODEX, CyCIF, IMC and MIBI), whereas the outer ring shows the corresponding disease composition within each technology. Bar plots summarize the disease compositions of the Melanoma dataset within IMC and of the MIBI technology, providing detailed proportions for these low-abundance categories, with diseases ordered from bottom to top by decreasing abundance. The color legend beneath the rings follows the same ordering and denotes the corresponding disease for each color.

Following pretraining, Spatium representations can be flexibly adapted to diverse downstream spatial proteomics applications through task-specific supervision (Fig. 2a). Rather than requiring separate models for individual analytical objectives, the pretrained encoder provides a shared representation space that can be optimized according to different biological questions. For cellular annotation, Spatium embeddings are coupled with a supervised classification head to transfer cellular identities from annotated reference datasets to previously unseen query datasets. This enables consistent cell type annotation across tissues and technologies while reducing dependence on extensive manual labeling. Beyond individual cell identities, Spatium captures higher-order tissue organization through spatial niche modeling. By predicting neighborhood cellular composition from learned cell representations, the model is encouraged to encode local multicellular context. The resulting embeddings can then be clustered to identify spatially coherent microenvironmental states, providing a representation-based approach for discovering tissue niches. Spatium also supports quantitative reconstruction of molecular states through protein imputation. Unlike cell-level prediction tasks that rely on the global cell embedding, protein imputation utilizes contextualized protein representations and incorporates complementary supervision from continuous protein abundance measurements and protein-specific abundance distributions. This enables recovery of missing protein information from incomplete spatial proteomics panels while preserving biologically meaningful expression patterns. Together, these task-specific adaptation strategies demonstrate that Spatium establishes a unified representation framework capable of supporting diverse spatial proteomics analyses, ranging from cellular annotation and tissue organization to molecular reconstruction.

### Few-Shot Cell Type Annotation from Sparse Spatial Proteomics

Spatial proteomics data are characterized by intrinsically low-dimensional and weakly informative protein measurements, where only a limited subset of markers exhibit strong and reproducible expression patterns across cell populations. Unlike high-dimensional transcriptomic profiles, the feature space is constrained and highly structured, leading to substantial ambiguity in cell identity inference under sparse observations.

Cell identities in spatial proteomics are typically characterized by combinatorial patterns of a limited set of canonical protein markers, consistent with marker-based cell phenotyping in multiplexed imaging and cytometry-based assays[17, 22]. In spatial proteomics datasets considered in this study, we further observe that the effective discriminative signal is concentrated in a small subset of markers, leading to a weak feature and effectively low-dimensional representation structure. This suggests that cell identity is encoded in a relatively compact and compositional marker signature space, rather than in high-dimensional feature richness.

To evaluate whether Spatium can learn such invariant identity representations, we formulate the task as a few-shot cell type annotation problem under limited supervision, where only a small fraction of spatial regions is used for training while the remaining regions are held out for evaluation within the same biological system. This design enforces that the model cannot rely on memorization of region-specific patterns but must instead infer cell identity from transferable marker combinations.

To evaluate the effectiveness of Spatium in this setting, we compare Spatium against a diverse set of representative baselines, including both general-purpose and domain-specific approaches. Specifically, we compare against: (i) linear probing and fine-tuning variants of Spatium embeddings, (ii) classical dimensionality reduction followed by linear regression (PCA), (iii) a cytometry-oriented variational autoencoder (CytoVI)[30], and (iv) Nicheformer[28], a foundation model primarily trained on imaging-based spatial data. These baselines are organized along three complementary axes: classical statistical representations (PCA), modality-adjacent representation learning methods (Nicheformer), and task-specific generative modeling for cytometry (CytoVI). This design enables a systematic comparison of representation quality under weak-feature, low-dimensional spatial proteomics settings. We evaluate performance under a few-shot cell type annotation setting, where the number of labeled samples per class is varied from 1 to 100. Across two independent spatial proteomics datasets generated from distinct imaging technologies (CODEX intestine dataset[31] and IMC CRC dataset[32]), comprising 57 and 14 protein panels, respectively (Fig. 3a) (Extended Data Fig. 1a), this setup evaluates model performance under extreme label scarcity across both platform and panel complexity.

**Fig. 3.**
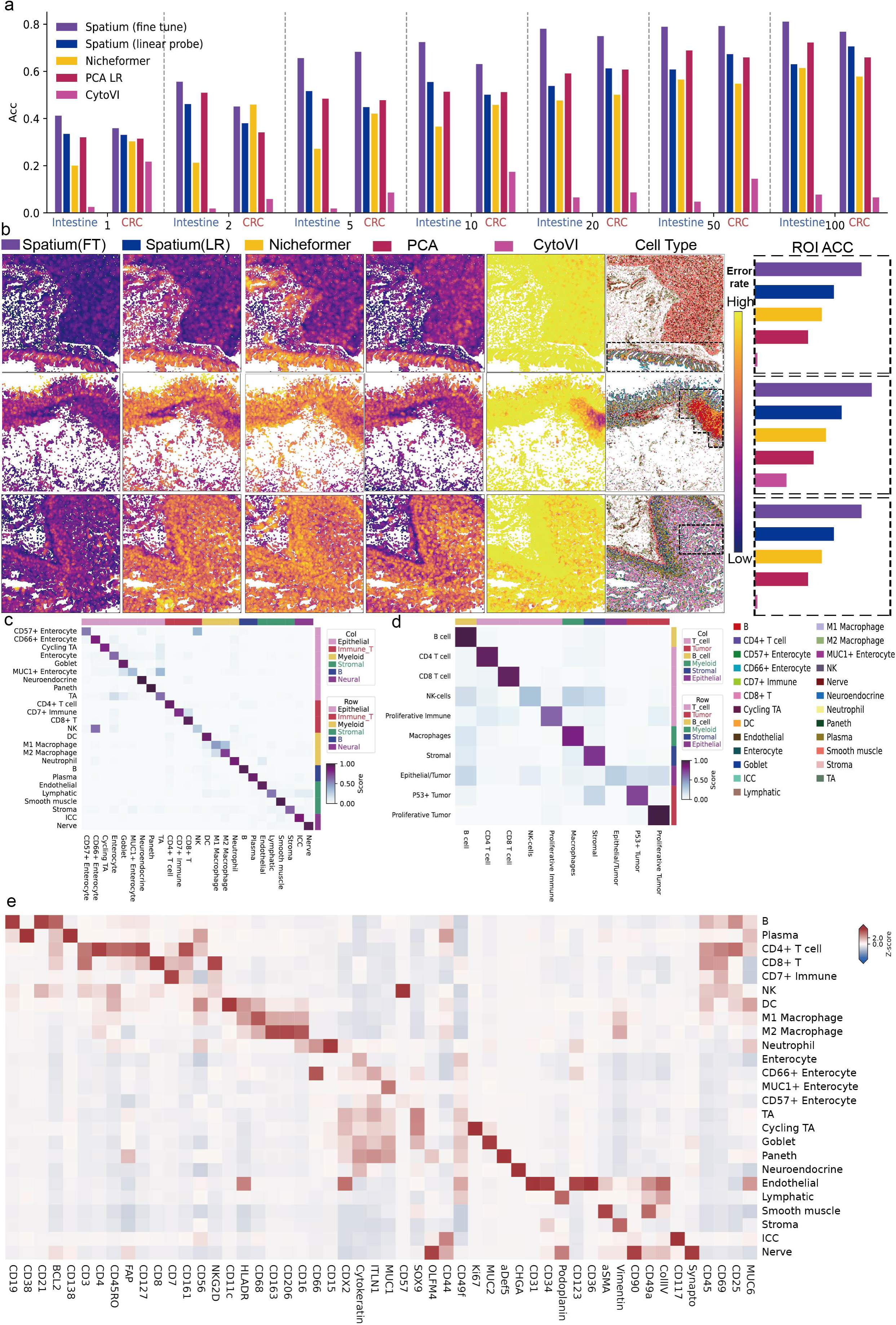
Few-shot spatial cell type annotation benchmarking of Spatium. (a). Benchmarking of Spatium and competing methods for few-shot spatial cell type annotation. Spatium was evaluated under both fine-tuning and linear probing settings and compared with Nicheformer, PCA-based linear regression (PCA LR), and CytoVI. For each dataset, reference sets were constructed by uniformly sampling 1, 2, 5, 10, 20, 50, or 100 cells per cell type, and the same reference/query splits were used across all methods. Classification performance was evaluated using accuracy (Acc) across different numbers of labeled cells. (b). Spatial visualization of annotation errors across representative regions of interest (ROIs). For each method, prediction errors were quantified by calculating the proportion of incorrectly annotated cells within the *k*-nearest-neighbor spatial neighborhood of each cell. Cells with higher local error rates are highlighted with increased intensity, indicating regions with higher annotation uncertainty. From top to bottom, the representative ROIs corresponds to the left colon, ileum, and mid jejunum. Error-prone spatial regions are outlined with dashed contours, and the corresponding annotation accuracies within each ROI are reported for comparison. (c, d). Confusion matrices of Spatium predictions on the intestine and CRC datasets, showing the distribution of correct and incorrect cell type assignments. Rows represent predicted cell types and columns represent ground-truth cell types. Color intensity indicates the proportion of cells assigned to each predicted class within the corresponding ground-truth cell type, with darker colors representing higher prediction frequencies. (e). Marker expression validation of Spatium predicted cell types in the intestine dataset.Heatmap shows average expression of marker genes across predicted cell types, with color intensity representing marker abundance.

As the number of labeled samples increases, all methods exhibit improved performance; however, Spatium maintains a clear and stable advantage across the entire scaling trajectory. In contrast, PCA-based linear models show competitive performance only in higher sampling case, while CytoVI remains unstable across sampling regimes, reflecting limited capacity to encode robust cell identity structure in low-dimensional protein space. Nicheformer shows improved performance as the number of reference cells increases but consistently underperforms Spatium, suggesting that representations learned from imaging-based spatial modalities do not fully transfer to spatial proteomics. Importantly, the gap between Spatium (fine-tuning) and Spatium (linear probe) highlights the benefit of task-adaptive refinement on top of strong pretrained representations, particularly in low-shot regimes where decision boundaries are highly sensitive to feature geometry.

To determine whether prediction errors occurred randomly or preferentially accumulated within specific tissue architectures, we next examined their spatial distribution across the query sections. We quantified the local density of misclassified cells by counting prediction errors within the neighborhood of each cell and visualized the resulting spatial error landscape (Fig. 3b) (Extended Data Fig. 1b, c).

Across all query regions, Spatium fine-tuning consistently exhibited the lowest spatial error density, with prediction errors remaining sparse and spatially isolated. In contrast, all baseline methods produced pronounced contiguous clusters of misclassifications, indicating reduced robustness within structurally complex tissue microenvironments. These error hotspots were consistently observed across multiple intestinal regions but varied substantially among different representation learning strategies.

Representative examples were observed in the Left colon and Ileum, where extensive prediction errors accumulated throughout complex mucosal compartments for nearly all baseline methods, whereas Spatium maintained accurate predictions across most of these regions. Notably, although PCA achieved relatively competitive overall classification accuracy, it still generated distinct error clusters within smooth muscle compartments, while Nicheformer exhibited systematic failures around ring-like smooth muscle structures surrounding endothelial cells. CytoVI produced widespread prediction errors across nearly all tissue architectures, reflecting its limited ability to learn discriminative representations for spatial proteomics data.

A similar pattern was observed in the Mid jejunum, where the tissue exhibits a layered organization spanning the mucosa and adjacent submucosa. Baseline methods accumulated large contiguous error regions across multiple tissue compartments, whereas Spatium largely preserved correct cell type assignments throughout these structurally heterogeneous regions, with only sparse local misclassifications.

Collectively, these observations demonstrate that the advantage of Spatium extends beyond improvements in overall classification accuracy. Rather than reducing the total number of prediction errors, Spatium substantially suppresses the formation of large, spatially coherent misclassification regions, indicating that Spatium learns cell identity representations that remain stable despite substantial local variation in marker composition and tissue architecture.

We next evaluated whether Spatium predictions preserve biologically coherent protein expression programs underlying cell identity. For each predicted cell type, we examined its dominant protein signals and assessed their consistency with canonical lineage-associated marker programs[31].

Across all major populations, Spatium produces highly structured and interpretable expression modules, where each cell type is characterized by a compact and specific set of proteins consistent with established biological definitions. These programs are clearly separated across major lineages, including epithelial, immune, and stromal compartments, indicating preservation of global cell identity organization in protein expression space.

Within epithelial populations, functional specialization is reflected by distinct marker programs, for example absorptive enterocytes enriched for cytokeratin, and secretory lineages such as goblet cells marked by MUC2[33]. Immune populations similarly exhibit coherent lineage-associated signatures, including CD4/CD8 T cell differentiation and macrophage-associated polarization markers (e.g., CD68, CD163)[34].

Overall, these results indicate that Spatium preserves the hierarchical organization of protein-defined cell identity, rather than producing entangled or over-smoothed representations, supporting its ability to recover biologically meaningful expression programs from spatial proteomics data.

### Spatium Captures Continuous Cell Identity Transitions in Ambiguous Protein States

Although Spatium achieved high overall annotation accuracy, we further investigated the biological characteristics of remaining prediction errors revealed by the confusion matrices (Fig. 3c, d). Rather than representing random classification failures, the major misclassification events were concentrated between biologically related cell populations with overlapping protein expression programs.

In the intestine dataset, the most prominent ambiguities involved NK cells and CD57+/CD66+ enterocyte populations. Specifically, 56.8% of annotated NK cells were predicted as CD66+ enterocytes, while 36.7% of CD57+ enterocytes were assigned to NK cells. To determine whether these errors reflected limitations of the model or intrinsic similarity in protein states, we compared the marker expression profiles of correctly classified and misclassified cells (Extended Data Fig. 2b, c).

For NK cells, correctly predicted NK populations exhibited marker patterns highly consistent with the original NK annotation, characterized by elevated expression of canonical immune markers including CD3, CD8, CD7, and CD21, together with strong CD57 expression (Extended Data Fig. 2b). In contrast, NK cells misclassified as CD66+ enterocytes showed a distinct shift in protein state, with reduced immune-associated marker signals and increased epithelial-associated signals, particularly CD66, resembling the expression profile of CD66+ enterocytes. These observations suggest that these cells represent an intermediate or shifted cellular state with attenuated NK-associated protein programs and partial acquisition of epithelial-like marker features, rather than indicating a systematic failure of the classifier.

A similar pattern was observed for CD57+ enterocytes misclassified as NK cells (Extended Data Fig. 2c). Correctly classified CD57+ enterocytes retained the expected epithelial signature characterized by high cytokeratin and CD57 expression, whereas misclassified cells displayed reduced epithelial marker expression and increased immune-associated signals, resulting in a protein profile closer to NK cells. Together, these results indicate that the observed confusion reflects continuous variation in protein-defined cell states rather than incorrect discrimination of well-separated cell identities.

To further validate whether these ambiguous predictions correspond to gradual shifts in protein identity states, we constructed simulation transition trajectories between NK and CD66+ enterocyte phenotypes by continuously interpolating their marker expression distributions. Using the original Spatium classifier without additional training, we evaluated how prediction probabilities changed along this simulated transition axis.

As the perturbation coefficient increased (Extended Data Fig. 2d), the simulated NK profiles gradually shifted toward CD66+ enterocyte-like states. Interestingly, the classification response followed a continuous transition rather than an abrupt switch. At low perturbation levels, cells were progressively reassigned from NK toward CD57+ enterocyte-like states, consistent with attenuation of NK-associated markers such as CD3, CD8, CD7, and CD21 while retaining partial immune-associated signals. With increasing perturbation, epithelial-associated features, including CD66 and cytokeratin, became dominant, resulting in a gradual increase in CD66+ enterocyte prediction probability. Notably, NK predictions disappeared at intermediate perturbation levels, while CD66+ enterocyte assignments became dominant only after stronger epithelial signature acquisition. This two-stage transition demonstrates that Spatium distinguishes gradual changes in marker composition and responds according to the underlying protein state landscape rather than relying on discrete memorized class boundaries. The gradual redistribution of prediction probabilities further indicates that Spatium learns a continuous representation of cell identity, allowing robust characterization of cells undergoing partial marker remodeling or transitional protein states.

### Neighborhood-aware embeddings enable biologically meaningful spatial niche identification

While cell-type annotation characterizes the identity of individual cells, spatial proteomics also seek to identify higher-order tissue niches defined by the composition of neighboring cells. Conventional neighborhood analysis[8] typically constructs neighborhood cell-type composition vectors and clusters these handcrafted features to identify recurrent microenvironmental states. We therefore asked whether the neighborhood-aware representations learned by Spatium inherently encode this spatial information.

To evaluate neighborhood encoding, we trained a regression head to predict the local neighborhood cell-type composition directly from the learned embeddings and compared the prediction performance with embeddings generated by PCA and CytoVI under the same regression framework. Spatium consistently achieved the highest cosine similarity between predicted and observed neighborhood composition across all ten major cell populations in CRC IMC dataset[32] (Fig. 4a), indicating that its representations more faithfully preserve local cellular organization than existing embedding methods.

**Fig 4.**
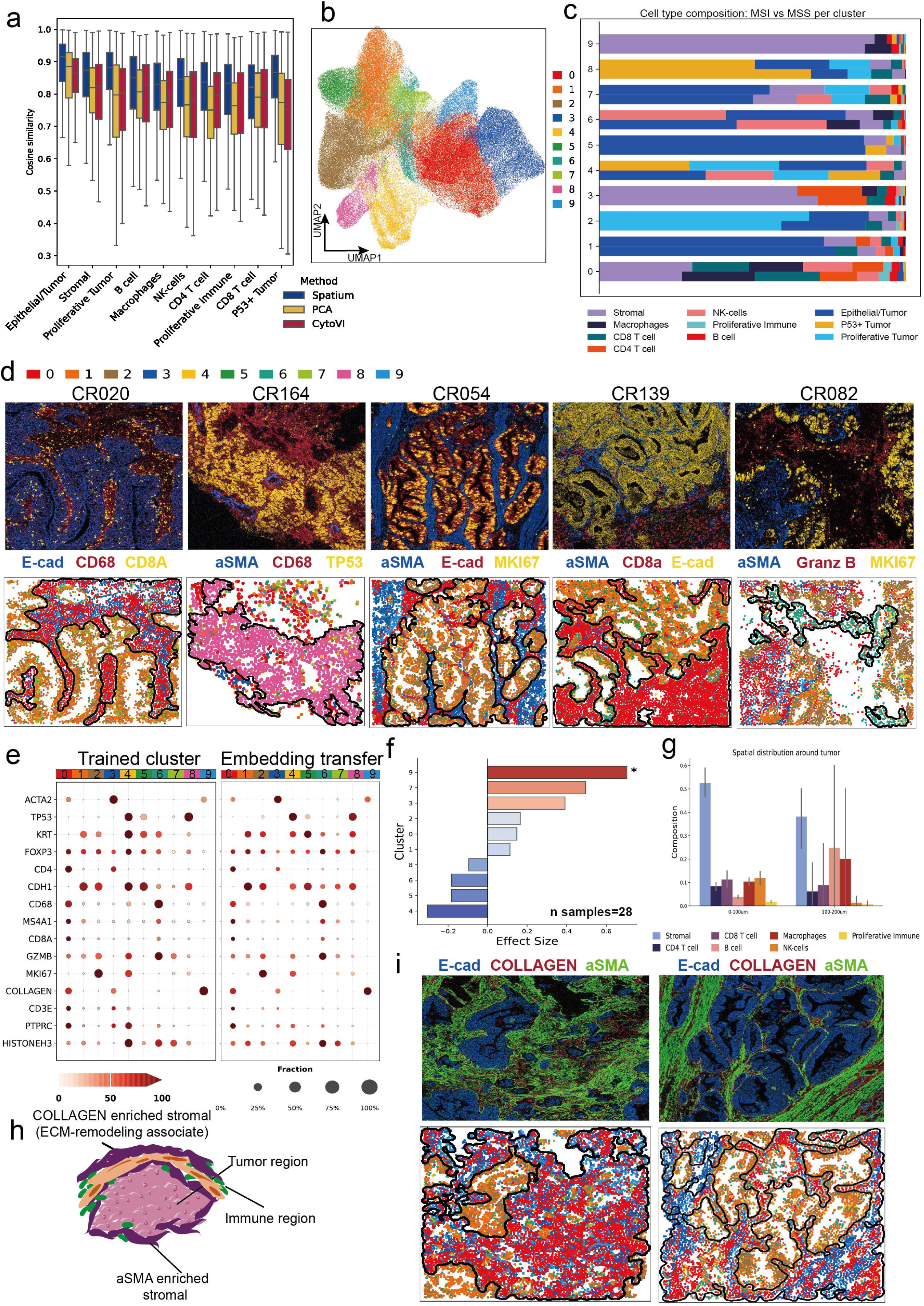
Identification and characterization of spatial niches using Spatium embeddings. (a). Evaluation of embedding quality for spatial niche identification. PCA, CytoVI, and Spatium derived embeddings were evaluated using a shared regression framework for autoregressive prediction of neighboring cellular compositions. Cosine similarity between predicted and observed cell-type compositions was calculated across ten cell types. (b). Identification of spatial niches based on Spatium embeddings. UMAP visualization of Spatium embeddings colored by unsupervised clustering assignments reveals ten distinct spatial niche clusters. (c). Cellular composition of spatial niche clusters across colorectal cancer (CRC) molecular subtypes. Stacked bar plots show the distribution of cell types within each spatial niche cluster across microsatellite instability (MSI) and microsatellite stable (MSS) tumors. For each cluster, the upper bar represents MSI samples and the lower bar represents MSS samples. Bar lengths indicate the relative proportion of each cell type within the corresponding niche cluster. (d). Spatial characterization of identified niches in representative CRC patients. The first row shows multiplexed imaging mass cytometry (IMC) images with selected markers highlighted according to their corresponding colors. The second row displays spatial niche assignments derived from Spatium embeddings. Dashed black contours indicate representative spatial regions corresponding to characteristic niche patterns in each patient. (e). Transferability of Spatium-derived spatial niche representations across independent CRC samples. Spatial niches were first identified by clustering Spatium embeddings from 10 clinically annotated training samples. For 19 additional samples without clustering-based niche annotations, niche labels were transferred from the reference samples based on embedding similarity. Dotplots show the average marker expression profiles of the original clustering-derived niches and transferred niches. (f). Association between spatial niches and CRC recurrence status. The relative abundance of each spatial niche cluster was quantified across 28 CRC samples and compared between recurrent and non-recurrent tumors. Statistical significance was assessed using the Mann–Whitney U test, and effect sizes were quantified using Cliff’s delta. Cluster 9 showed the strongest association with recurrence status, exhibiting the largest effect size and significant enrichment between groups with p-value= 0.01278. (g). Spatial distribution of major cell types surrounding tumor regions. The relative abundance of each cell type was quantified within concentric spatial regions located 0–100 μm and 100–200 μm from tumor boundaries. (h). Schematic illustration of representative spatial cellular organization patterns captured by Spatium-defined spatial niches. (i). Spatial visualization of characteristic spatial niches in representative CRC patients. The upper panels show IMC images with selected markers indicated by corresponding colors. The lower panels show spatial niche assignments generated from Spatium embeddings, with black contours highlighting the spatial distribution of niche 3 and niche 9.

We next clustered the learned neighborhood embeddings to identify recurrent spatial niches within the CRC IMC dataset. The resulting embedding space exhibited well-organized continuous manifolds, with closely related niches occupying neighboring regions while remaining clearly distinguishable from transcriptionally and spatially distinct microenvironments (Fig. 4b), suggesting that the learned representations capture biologically meaningful gradients of tissue organization.

Characterization of neighborhood composition together with marker enrichment enabled biological annotation of the identified spatial niches (Fig 4c, Extended Data Fig. 3a, b). Several niches represented distinct stromal states despite sharing fibroblast dominance. Cluster 3 was characterized by prominent ACTA2 enrichment together with substantial immune cell components, consistent with an activated CAF-associated niche[35, 36]. In contrast, cluster 9 exhibited the strongest collagen enrichment across all identified niches, accompanied by only moderate ACTA2 expression, defining a collagen-rich stromal niche potentially associated with extracellular matrix remodeling rather than simply increased fibroblast abundance[37]. Among epithelial niches, clusters 1 and 5 were enriched for epithelial and tumor cells and displayed strong CDH1 and KRT expressions, whereas cluster 2 represented a proliferative tumor niche distinguished by dominant proliferative tumor composition together with marked MKI67 enrichment. Immune-associated cluster 6 was characterized by enrichment of macrophages and NK cells, accompanied by elevated CD68 and GZMB expression. Together, these findings demonstrate that the learned neighborhood embeddings resolve biologically meaningful tissue microenvironments defined by coordinated cellular composition and molecular features rather than coarse cell-type abundance alone.

To examine whether the identified niches reflected genuine tissue organization, we visualized representative niche distributions across multiple CRC specimens and compared their spatial patterns with the corresponding multiplex immunofluorescence images (Fig. 4d, Extended Data Fig. 3c). For each specimen, characteristic niche regions were delineated by contour overlays, allowing direct comparison with the underlying protein-defined tissue architecture. Across all representative cases, niche boundaries closely recapitulated the spatial organization revealed by the original IMC signals. For example, in CR020, the immune-enriched niche (cluster 0) precisely outlined the CD68- and CD8A-positive immune structures surrounding E-cadherin (CDH1)-positive epithelial tumor regions. In CR054, the proliferative tumor niche spatially coincided with MKI67-positive tumor regions, while ACTA2-positive stromal fibers corresponded to the activated stromal niche. Likewise, TP53-enriched tumor regions (CR164) and GZMB-positive NK-rich regions (CR082) were accurately captured by their corresponding niche distributions. These observations demonstrate that the identified niches preserve higher-order spatial tissue architecture rather than representing abstract clusters in embedding space.

### Transferable spatial niches reveal a recurrence-associated ECM-associate microenvironment

To examine whether the learned niche representations generalize across independent specimens, we transferred the niche labels learned from the training cohort to an additional cohort using embedding similarity without re-clustering. Despite being assigned entirely through embedding-based label transfer, transferred niches preserved nearly identical marker enrichment patterns to those observed in the original training cohort (Fig. 4e), demonstrating that the learned neighborhood representations capture reproducible biological states rather than dataset-specific clustering structures.

Having established the robustness of niche transfer, we next investigated whether specific spatial niches were associated with clinical outcome. Comparing niche abundance between recurrent and non-recurrent CRC patients identified the collagen-rich ECM-remodeling associate niche as the only significantly enriched niche in recurrent tumors, exhibiting both the largest effect size and the strongest enrichment among all identified niches[38] (Fig. 4f). In contrast, other stromal niches, including the ACTA2-enriched activated CAF niche[39], showed substantially weaker associations with recurrence. To further characterize the spatial organization of stromal niches, we examined their distribution relative to tumor regions (Fig. 4g–h). Activated CAF-associated niches predominantly formed extensive stromal networks surrounding tumor regions, whereas the ECM-remodeling niche consistently appeared as compact collagen-enriched substructures embedded within these larger stromal architectures. This hierarchical spatial organization was consistently observed across multiple CRC specimens.

Representative tissue sections further illustrate this structural relationship (Fig. 4i). Immunofluorescence staining showed that ACTA2-positive stromal networks closely matched the distribution of activated CAF-associated niches, while regions enriched for collagen corresponded specifically to the ECM-remodeling niche nested within these stromal structures. These observations indicate that the ECM-remodeling niche represents a specialized stromal state characterized by localized extracellular matrix remodeling rather than a simple increase in fibroblast abundance[37].

Collectively, these analyses identify a reproducible collagen-rich ECM-remodeling niche that can be robustly transferred across independent cohorts and is preferentially enriched in recurrent colorectal cancer. The spatial organization and molecular characteristics of this niche suggest that localized extracellular matrix remodeling represents a distinct stromal state associated with recurrence-associated tumor ecosystems.

### Spatium generalizes protein imputation across spatial proteomics datasets

Beyond cell annotation and spatial niche characterization, recovering unmeasured proteins represents another important downstream application in spatial proteomics. Because different imaging panels frequently profile only a subset of proteins, accurate imputation of missing markers could facilitate biological interpretation and improve the utility of partially observed datasets. To evaluate whether the protein-state representations learned by Spatium support this task, we developed a protein imputation framework in which selected proteins were masked from an independent validation section and reconstructed from the remaining measured proteins. For each dataset, the prediction head was trained using five tissue sections and evaluated on a held-out section, allowing direct comparison between predicted and experimentally measured protein expression without information leakage.

Representative results from the HCC CODEX dataset[40] demonstrate that Spatium accurately reconstructs the spatial distributions of multiple masked proteins (Fig. 5a,c). Predicted rank-normalized expression patterns closely matched the experimentally measured distributions across individual cells while preserving their corresponding tissue organization. For example, reconstructed Pan-cytokeratin faithfully localized to biliary epithelial regions, CD8 coincided with infiltrating CD8 T-cell populations, and Podoplanin accurately delineated lymphatic endothelial structures, demonstrating that the learned representations preserve biologically meaningful protein-state information rather than simply reproducing global expression trends.

**Fig 5.**
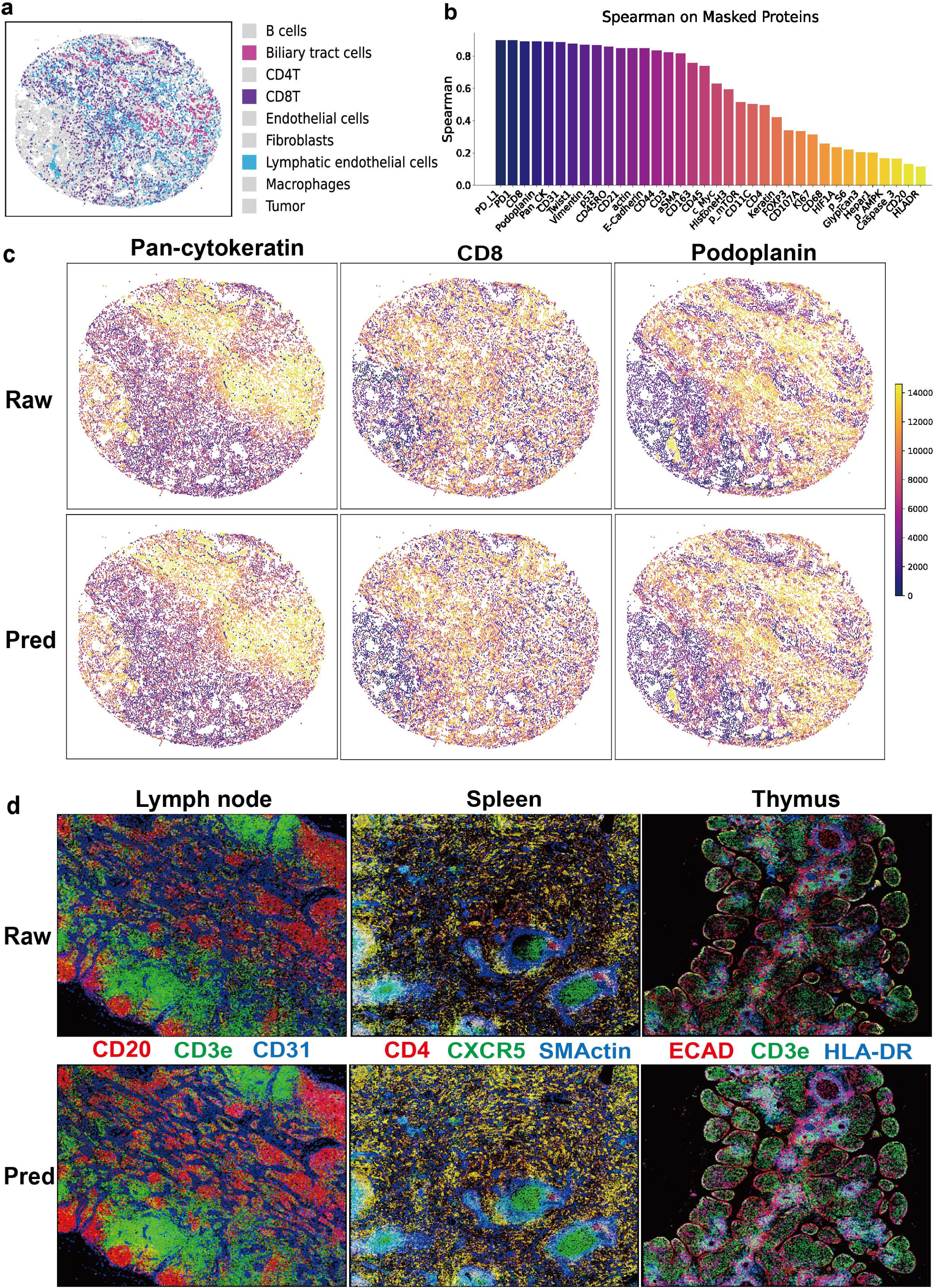
Spatial protein imputation using Spatium embeddings. (a). Spatial reconstruction of masked proteins in the HCC CODEX dataset. Representative spatial distributions of three major cell populations, including biliary tract cells, CD8 T cells, and lymphatic endothelial cells, are shown in the validation sample. Protein markers were masked from the input data and reconstructed using the Spatium fine-tuning framework. (b). Quantitative evaluation of protein imputation performance in the HCC CODEX dataset. For each protein, expression values were masked in the validation sample and reconstructed by Spatium. Spearman correlation coefficients between the predicted and measured protein abundances are shown for individual proteins. (c). Comparison of measured and imputed protein expression patterns at the single-cell level. For representative proteins, including Pan-cytokeratin, CD8, and Podoplanin, measured and predicted expression values were converted into within-sample ranks to enable comparison of relative protein abundance across cells. (d). Fluorescence image reconstruction of masked proteins across three CODEX datasets, including HuBMAP CODEX lymph node, spleen CODEX, and thymus CODEX. For each dataset, the upper row shows the reconstructed fluorescence intensity images generated from imputed protein expression values, and the lower row shows the corresponding original CODEX fluorescence images. Representative masked protein channels are displayed to compare the predicted and measured fluorescence intensity patterns at the same cellular locations.

Quantitative evaluation further confirmed robust reconstruction performance across a broad range of proteins (Fig. 5b). Lineage-associated and structural markers, including Pan-Cytokeratin, Podoplanin, CD8, CD31, PD-L1, PD1, Twist1, Vimentin and p53, achieved Spearman correlations approaching or exceeding 0.85, indicating highly accurate recovery of their spatial expression patterns. Although several signaling-related proteins exhibited lower prediction accuracy, likely reflecting their greater context dependence and intrinsic biological variability, the overall results demonstrate that Spatium embeddings retain sufficient information to reconstruct diverse protein identities from incomplete measurement panels.

To further evaluate whether reconstructed proteins preserve higher-order tissue architecture, we compared predicted protein maps with the original immunofluorescence images across multiple independent tissues (Fig. 5d, Extended Data Fig. 4b, c, d). In lymph node, reconstructed CD20, CD3E and CD31 faithfully recovered the characteristic organization of B-cell follicles, T-cell zones and vascular structures. Similar spatial consistency was observed in spleen, where predicted CD4, CXCR5 and smooth muscle actin (ACTA2) reproduced the compartmental organization surrounding splenic vascular structures. Likewise, reconstructed E-cadherin (CDH1), CD3E and HLA-DR in thymus accurately recapitulated the characteristic epithelial and lymphoid architectures observed in the measured images. Across all examined tissues, reconstructed proteins preserved coherent spatial organization and tissue compartment boundaries, indicating that Spatium recovers biologically meaningful protein landscapes rather than isolated per-cell expression values.

Additional evaluations across the HuBMAP lymph node, spleen and thymus datasets[41, 42] further demonstrated the generalizability of protein reconstruction (Extended Data Fig. 4a). Reconstruction accuracy gradually decreased as progressively larger fractions of proteins were masked, exhibiting graceful performance degradation despite increasingly limited input information. Furthermore, randomly permuting the within-cell protein ordering before inference almost completely abolished prediction accuracy, confirming that successful reconstruction depends on the learned protein-state representations rather than dataset-specific priors or individual marker abundances. Together, these results establish protein imputation as a practical downstream application of Spatium and demonstrate that the learned representations capture sufficient biological information to infer missing proteins while preserving spatial tissue organization.

### Spatium enables unsupervised discovery of disease-associated cellular states and spatial hematopoietic niches

A key property of foundation models is the ability to capture transferable biological representations beyond explicitly supervised tasks. To assess whether Spatium embeddings preserve biologically meaningful cellular states without task-specific labels, we applied Spatium to a CODEX spatial proteomics cohort consisting of five acute myeloid leukemia (AML) bone marrow samples and three normal bone marrow (NSM) samples[43]. Rather than using cell type annotations or disease labels, we adapted Spatium only through the masked language modeling objective and extracted cell-level representations for downstream analysis (Fig. 6a, Extended Data Fig. 5a).

**Fig 6.**
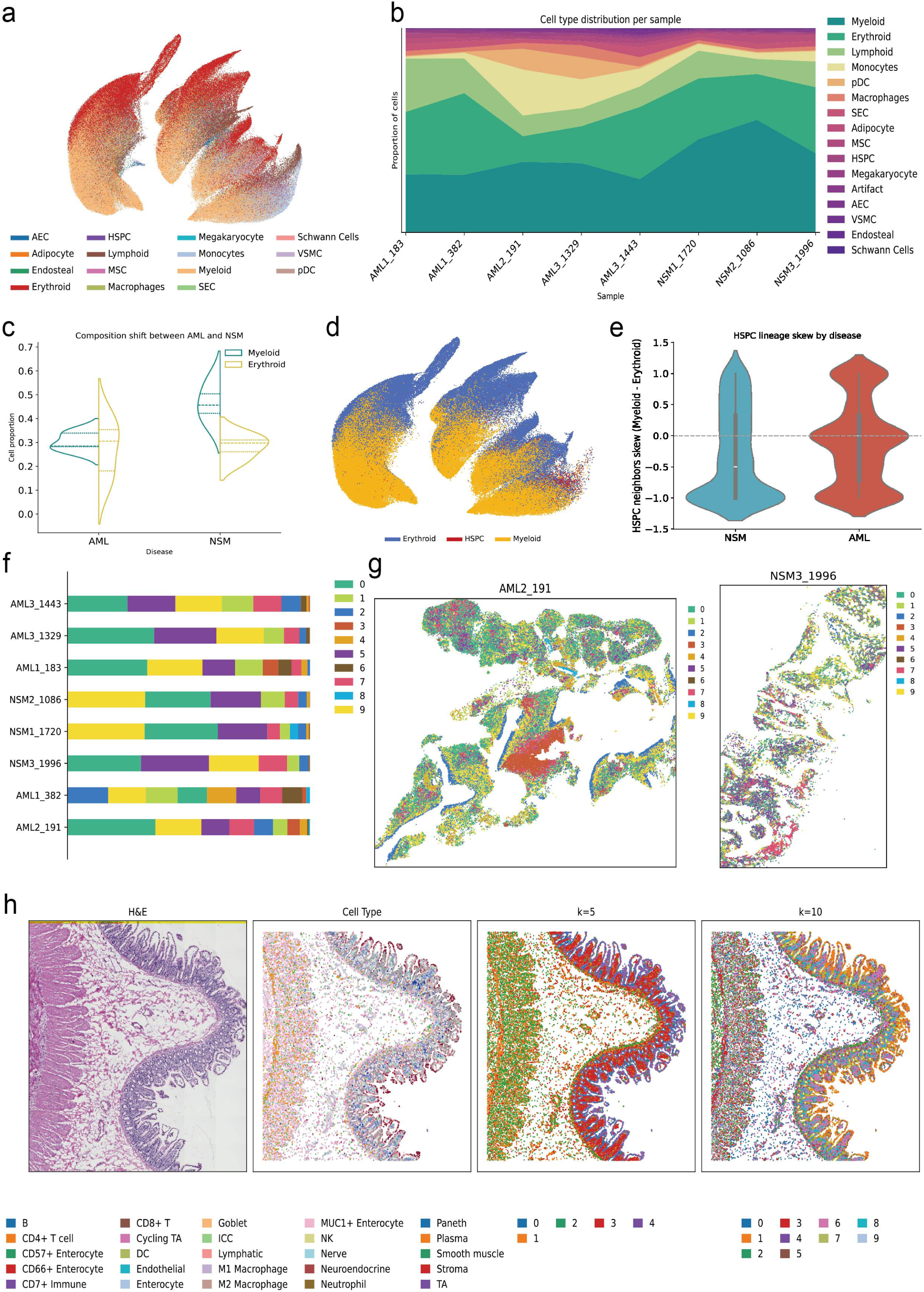
Label-free analysis of bone marrow CODEX data using Spatium embeddings. (a). Unsupervised characterization of human bone marrow CODEX samples using Spatium embeddings. Eight bone marrow samples, including five acute myeloid leukemia (AML) samples and three non-malignant samples (NSM), were fine-tuned using the masked language modeling (MLM) objective without cell type labels. UMAP visualization of the resulting Spatium embeddings is shown, with cells colored by reference cell type annotations. (b). Cell type composition across human bone marrow CODEX samples. The relative abundance of annotated cell types is shown for each AML and NSM sample. (c). Comparison of Myeloid and Erythroid cell proportions between AML and NSM samples. The relative abundance of Myeloid and Erythroid populations is compared between the two groups, with each point representing an individual sample. (d). Identification and visualization of HSPC populations within Spatium embeddings. UMAP plots show the distribution of HSPC cells across AML and NSM samples, with highlighted HSPC populations displayed together with neighboring cellular states. (e). Neighborhood composition analysis of HSPC cells. The relative abundance of Myeloid and Erythroid populations within the local spatial neighborhoods surrounding HSPC cells is quantified across AML and NSM samples. (f). Spatial niche composition across human bone marrow CODEX samples. The relative abundance of identified spatial niches is shown for each sample. (g). Spatial visualization of representative bone marrow niches. Representative CODEX images and corresponding spatial niche assignments are shown for selected samples, illustrating the spatial organization of identified niches. (h). Integration of histology-derived and spatial protein embeddings. HE images from human intestine samples were processed using a histology foundation model to generate cell-level image embeddings, which were integrated with Spatium embeddings. Visualization shows the original HE images, reference cell type distributions, and clustering results from the integrated embeddings using different neighborhood parameters (*k* = 5 and *k* = 10).

Despite the absence of supervised information, Spatium embeddings recovered structured cellular landscapes across individual bone marrow samples. UMAP visualization revealed continuous cellular state distributions rather than isolated clusters, with comparable organization observed across different samples (Fig. 6a). These results indicate that the pretrained representation space preserves intrinsic biological variation and can capture cellular state relationships without explicit annotation guidance. We next investigated whether these unsupervised representations reflected disease-associated remodeling of hematopoietic composition. Comparison of cellular populations between AML and NSM samples revealed distinct alterations in hematopoietic organization (Fig. 6b). NSM samples exhibited increased proportions of erythroid and myeloid populations, whereas AML samples showed reduced myeloid representation with substantial inter-sample variability (Fig. 6c). These observations demonstrate that Spatium embeddings can recover disease-associated differences in cellular composition without prior knowledge of disease status.

To further explore the cellular states associated with these differences, we examined hematopoietic stem and progenitor cell (HSPC)-associated populations within the embedding space. HSPC populations were markedly expanded in AML samples compared with NSM controls and occupied intermediate regions between myeloid- and erythroid-associated cellular states[44] (Fig. 6d). This positioning suggests that Spatium embeddings capture continuous hematopoietic state transitions rather than representing cell identities as isolated categories.

We next investigated the spatial context surrounding HSPCs by quantifying the cellular composition of their local neighborhoods. HSPC-associated neighborhoods displayed distinct cellular biases between AML and NSM samples (Fig. 6e). In NSM samples, HSPCs were preferentially surrounded by myeloid-dominant neighborhoods, whereas AML samples exhibited altered local cellular compositions, suggesting disease-associated remodeling of HSPC spatial environments. We next investigated the spatial context surrounding HSPCs by quantifying the cellular composition of their local neighborhoods. HSPC-associated neighborhoods displayed distinct cellular biases between AML and NSM samples (Fig. 6e). In NSM samples, HSPCs were preferentially surrounded by myeloid-dominant neighborhoods, whereas AML samples exhibited altered local cellular compositions, suggesting disease-associated remodeling of HSPC spatial environments.

To further characterize higher-order spatial organization, we applied the Spatium neighborhood representation framework to identify spatial niches (Fig. 6f, g, Extended Data Fig. 5b). Several niche populations showed strong enrichment of HSPCs, with clusters 0, 5 and 7 exhibiting the highest HSPC enrichment scores (Extended Data Fig. 5c). These HSPC-associated niches displayed distinct cellular compositions, including myeloid-dominant, erythroid-dominant and lymphoid-associated environments, respectively (Extended Data Fig. 5d). Importantly, spatial visualization confirmed that these niches formed coherent tissue structures rather than random distributions, demonstrating that Spatium representations capture organized hematopoietic microenvironments (Fig. 6g).

We next examined the molecular characteristics of HSPC-associated states through marker enrichment analysis of rare HSPC-enriched embedding clusters. Several rare populations consistently exhibited elevated SPINK2 expression together with CD138, BCL2, CD44 and oxidative phosphorylation-associated signatures (Extended Data Fig. 5e)[45, 46]. Although these populations represent a small fraction of total cells, their reproducible molecular patterns suggest that Spatium can resolve specialized cellular states within heterogeneous hematopoietic environments that may be overlooked by conventional annotation strategies. Notably, these molecularly distinct HSPC states were identified within AML-associated spatial niches, suggesting that disease-associated microenvironmental remodeling is accompanied by the emergence of specialized HSPC populations with primitive, survival-associated, and niche-interactive features[44]. Together, these results demonstrate that Spatium embeddings capture biologically meaningful cellular structure and spatially organized microenvironments without requiring supervised labels. Beyond predefined annotation tasks, Spatium enables unsupervised discovery of latent cellular states and disease-associated spatial organization in complex tissues.

### Multimodal transfer and integration through Spatium representations

Spatial proteomics provides direct access to protein-level cellular states, but complementary molecular and imaging modalities contain additional information that is not directly captured by protein marker panels. We therefore explored whether Spatium representations could serve as a unified latent space for transferring and integrating information across heterogeneous spatial biology modalities.

To evaluate whether Spatium could bridge transcriptomic and proteomic measurements, we first investigated cross-modal cell-type inference between annotated scRNA-seq references and CODEX spatial proteomics datasets. Protein markers measured in CODEX were mapped to their corresponding gene symbols, and scRNA-seq profiles were converted into Spatium-compatible protein token sequences using the same unified vocabulary used during spatial proteomics pretraining. The pretrained Spatium backbone was subsequently fine-tuned using annotated scRNA-seq cells, and the learned classifier was directly applied to spatial proteomic cells without additional adaptation. In human tonsil CODEX data[19, 47, 48], this cross-modal transfer strategy achieved an overall cell-type annotation accuracy of 0.7869, demonstrating that transcriptomic information could be effectively projected into the spatial proteomic representation space (Supplementary Fig. 1). Predicted cell populations showed strong correspondence with their expected protein marker profiles (Supplementary Fig. 2). For example, CD4 T and CD8 T populations exhibited elevated CD4 and CD8 expression patterns, respectively, whereas B-cell populations were characterized by increased B-lineage markers including CD19, CD22 and CD40. Plasma-like populations showed enrichment of CD38 and CD63-associated signatures, while dendritic cell predictions exhibited elevated CD123 and CD1c expression. Together with the confusion matrix analysis, these results demonstrate that Spatium captures modality-independent cellular identity features that enable transfer of annotations across transcriptomic and protein-based measurements.

Beyond molecular transfer, we further investigated whether Spatium representations could incorporate complementary morphological information from histological imaging. Paired H&E images from intestinal spatial proteomics datasets were processed using the pretrained pathology foundation model Prov-GigaPath[49] to obtain cell-level image representations. These histological embeddings were then aligned with Spatium protein embeddings through a learnable fusion framework, generating joint multimodal representations.

The resulting fused embeddings preserved both tissue-level organization and cellular heterogeneity captured by spatial proteomics (Fig. 6h). Compared with the original H&E morphology and protein-derived cell-type distributions, multimodal embeddings recovered spatial structures consistent with the underlying tissue architecture while simultaneously refining cell-level organization. At different clustering resolutions, the fused representations revealed hierarchical tissue patterns, where coarse clusters corresponded to major anatomical structures and finer clustering resolved cellular substructures embedded within these regions. These results suggest that Spatium representations can be extended beyond protein-only measurements and provide a flexible framework for integrating spatial proteomics with complementary molecular and imaging modalities.

## Discussion

Spatial proteomics has emerged as a powerful approach for resolving cellular organization and molecular states within intact tissues, yet the field currently lacks a general representation framework capable of learning transferable biological features across diverse experimental settings. Existing computational approaches are often optimized for individual datasets, rely heavily on predefined marker panels or manual annotation strategies, and are difficult to generalize across technologies with different measurement characteristics. Here, we introduce Spatium, a foundation model for spatial proteomics that learns transferable representations of cellular protein states from 51 million cells across 70 spatial proteomics studies. By converting heterogeneous protein measurements into a unified ranked representation space, Spatium enables a shared computational framework for diverse downstream analyses, including cell-type annotation, spatial niche discovery, protein state reconstruction and multimodal integration.

A central challenge in spatial proteomics is the substantial heterogeneity introduced by experimental design. Unlike transcriptomic assays, where measurements are typically performed within a relatively stable genomic feature space, spatial proteomic platforms rely on curated antibody panels that vary substantially across studies, tissues and technologies. Furthermore, measured fluorescence intensities are affected by platform-specific characteristics, antibody performance and tissue context, limiting direct comparison of absolute protein abundance across datasets. To address these challenges, Spatium adopts a rank-based protein tokenization strategy that represents cells according to relative protein abundance patterns rather than absolute intensity values. This design allows the model to focus on combinatorial protein states that define cellular identity while reducing sensitivity to study-specific measurement variation. The success of this strategy highlights the potential of relative molecular organization as a robust representation principle for heterogeneous spatial proteomic data.

The learned Spatium representations capture transferable cellular identity information across tissues, technologies and annotation scenarios. Through few-shot cell-type annotation experiments, Spatium maintained strong performance even when only limited labeled examples were available, demonstrating that large-scale pretraining can reduce dependence on extensive manual annotation. Such transferability is particularly important for spatial proteomics, where clinical samples and rare tissues frequently lack comprehensive reference annotations. Rather than requiring dataset-specific marker selection and handcrafted rules, Spatium provides a reusable representation space that enables cellular states learned from large-scale datasets to be applied to previously unseen spatial experiments.

Beyond individual cell identities, Spatium extends representation learning toward higher-order tissue organization. Conventional spatial niche analysis often relies on manually defined neighborhood compositions followed by clustering of cell-type proportions, which captures local cellular abundance but provides limited access to underlying biological states. By training representations to encode neighborhood composition and extracting contextualized embeddings, Spatium enables discovery of spatially organized microenvironmental states that reflect complex combinations of cellular populations and molecular programs. These learned representations further supported unsupervised exploration of disease-associated cellular states in human bone marrow, where Spatium identified rare HSPC-associated populations and spatial niches associated with altered hematopoietic organization. Together, these findings demonstrate that foundation models can provide a route toward unbiased discovery of latent tissue states beyond predefined annotation frameworks.

In addition to representation-based discovery, Spatium demonstrates the potential to recover missing molecular information from incomplete spatial proteomic measurements. Spatial proteomic experiments are frequently constrained by limited marker panels, restricting the number of biological features that can be simultaneously measured. By leveraging contextual relationships among protein states learned during pretraining, Spatium can reconstruct unmeasured protein patterns while preserving spatial distributions and protein-specific abundance characteristics. Although current evaluations focus primarily on within-dataset reconstruction, this capability provides a foundation for future development of cross-panel and cross-platform protein imputation approaches, which may expand the biological coverage of existing spatial proteomic experiments.

The ability of Spatium to integrate information across modalities further suggests that spatial proteomics foundation models may serve as general representation layers for spatial biology. By transferring cellular identities from scRNA-seq references to spatial proteomic datasets and integrating histological image representations with protein-derived embeddings, Spatium demonstrates that learned protein representations can relate to complementary molecular and morphological information. Such multimodal integration may become increasingly important as spatial technologies continue to evolve, enabling joint analysis of molecular measurements, tissue morphology and other spatially resolved biological signals within a common computational framework. Despite these advances, several limitations remain. First, spatial proteomic measurements are inherently constrained by the available antibody panels and do not yet represent comprehensive proteome-level information. Future extensions incorporating larger marker panels, additional protein measurements or complementary omics modalities may further improve the breadth of learned representations. Second, although Spatium learns cellular representations and incorporates neighborhood information in downstream analyses, the current pretraining strategy primarily models cell-level protein states. Incorporating explicit spatial architectures, such as graph-based or spatial attention mechanisms, may enable direct learning of tissue topology during pretraining. Third, protein imputation and multimodal integration require further validation across independent technologies and experimental settings to fully assess their generalizability. Finally, while Spatium reveals disease-associated spatial states, larger clinical cohorts and functional studies will be required to determine how these computationally identified states contribute to disease progression and therapeutic response.

In summary, Spatium establishes a foundation model framework for spatial proteomics by learning transferable representations from large-scale heterogeneous datasets. By enabling cross-dataset annotation transfer, spatial niche discovery, molecular state reconstruction and multimodal integration, Spatium provides a general computational foundation for extracting biological insights from increasingly diverse spatial proteomic technologies. As spatial measurements continue to expand in scale and complexity, representation learning approaches such as Spatium may facilitate the transition of spatial proteomics from dataset-specific analysis toward a more unified and scalable computational paradigm.

## Methods

### Spatium datasets

#### Spatium pretrain datasets

The pretraining dataset for Spatium comprises four spatial proteomics technologies, including CODEX, Imaging Mass Cytometry (IMC), Cyclic Immunofluorescence (CyCIF), and Multiplexed Ion Beam Imaging (MIBI). For CODEX data, we collected publicly available datasets from multiple resources, including the HuBMAP portal[41, 42], scProAtlas[50], Aquila[51], and other curated studies providing processed and downloadable expression matrices. For IMC, CyCIF, and MIBI, datasets were similarly obtained from previously published studies as well as processed files provided by scProAtlas.

In total, all data were organized into 70 datasets stratified by study source and tissue type. Given substantial heterogeneity across studies in both file formats and protein panel nomenclature, we performed a unified preprocessing procedure prior to model training. Specifically, protein panel names were standardized by converting all markers to uppercase and mapping them to a unified global protein vocabulary. After harmonization, each dataset was independently structured into an AnnData object[52] to ensure consistency in downstream processing and model input formatting.

#### Human intestine dataset

The human intestine CODEX dataset[31] consists of eight regions of interest (ROIs) derived from two healthy human samples, covering multiple anatomical sites of the gastrointestinal tract. Specifically, the Duodenum, Proximal jejunum, Right colon, and Transverse colon were used to construct the support set, while the Left colon, Ileum, Mid jejunum, and Sigmoid colon were designated as the query set. All 25 annotated cell types and the original data preprocessing pipeline were obtained directly from Hickey et al.[31], which served as the reference for downstream model evaluation. Both support and query sets share a common protein panel consisting of 57 markers. The query set contains 214,837 cells and is fully utilized for inference and performance evaluation. The support set contains 184,600 cells, from which we perform balanced subsampling across cell types. Specifically, we uniformly sample *k* cells per cell type, where *k* is set to 1, 2, 5, 10, 20, 50, and 100, respectively, to evaluate model performance under varying levels of support data availability. All processed CODEX data can be accessed from original paper.

#### CRC dataset

The IMC colorectal cancer (CRC) dataset consists of 52 tissue samples spanning 162 tissue regions, accompanied by clinical metadata, raw IMC images, processed IMC expression matrices, and annotations of 10 cell types provided by Su et al.[32]

For our study, we selected 28 cohorts with complete clinical information to ensure consistency across samples. The resulting dataset contains 14 protein markers and a total of 457,854 cells. or few-shot learning evaluation, we designated CR144 as the support set (7,629 cells) and CR101 as the query set (19,964 cells), enabling controlled assessment under a fixed support–query split. These two cohorts were used to assess model performance under controlled support-query separation. For embedding-based clustering and niche label transfer analysis, all 28 cohorts were jointly utilized to enable cross-sample representation learning and spatial domain alignment. All raw IMC data, processed expression matrices, and corresponding clinical annotations were obtained directly from the original publication without modification. The 14-marker panel covers major cellular compartments, including tumor, stromal, epithelial, and immune markers. Detailed descriptions of each marker and its associated biological function are provided in Supplementary Table 1.

#### HCC dataset

The CODEX hepatocellular carcinoma (HCC) dataset was derived from the large-scale CODEX study reported by Qiu et al.[40], which originally comprised 401 HCC tissue samples collected in a tissue microarray (TMA) cohort. Each CODEX region represents distinct tumor microenvironments with varying compositions of tumor, immune, and stromal cells. For the protein imputation task, we constructed the training dataset using five representative tissue regions, resulting in a training set containing 63,849 cells with complete measurements of 33 protein markers. An independent tissue region was used as the validation set, comprising 14,607 cells with the same 33-marker panel. To evaluate protein imputation performance, we randomly removed *p* protein markers from the validation set, where the selected markers were completely excluded from all cells. During the ablation experiments, *p* was gradually increased from 5 to 33, resulting in progressively more challenging imputation tasks. The 33 protein markers collectively cover tumor, immune, and stromal cell populations.

#### Human lymph node, spleen, thymus dataset

The CODEX datasets of human lymph node, spleen, and thymus tissues were obtained from the HuBMAP portal (https://portal.hubmapconsortium.org/)[41, 42]. For the protein imputation task, we constructed dataset-specific training and validation splits for each tissue type. The lymph node, spleen, and thymus training sets contain 497,986, 579,440, and 311,256 cells, respectively, with 29, 24, and 28 protein markers. The corresponding validation sets contain 90,504, 147,676, and 102,780 cells, respectively, measured over the same marker panels. The imputation training and evaluation protocol follows the same setting as in the HCC dataset, where missing proteins are simulated in the validation set by progressively removing protein markers. All raw fluorescence images and corresponding segmentation masks were downloaded from HuBMAP using the respective tissue identifiers. The processed CODEX expression matrices were provided in AnnData format for downstream analysis.

#### Human bone marrow dataset

The human bone marrow CODEX dataset was obtained from the study by Bandyopadhyay et al.[43], which includes eight tissue sections spanning distinct anatomical regions and disease conditions. Specifically, the dataset consists of five acute myeloid leukemia (AML) samples and three negative lymphoma staging bone marrow biopsies (NSM), covering diverse pathological states of the bone marrow microenvironment. The full dataset contains 327,563 cells profiled over a 50-marker protein panel. All cell-type annotations were adopted from the original study without modification. For downstream analysis, including embedding learning and clustering, all eight tissue sections were jointly used for model training and inference to enable cross-sample representation learning.

#### Human tonsil dataset

The human tonsil CODEX and matched scRNA-seq datasets were obtained from the MaxFuse[19, 47, 48] study. The CODEX dataset contains 178,919 cells profiled with a 47-marker protein panel, while the matched scRNA-seq dataset contains 12,977 cells and 33,538 genes. Cell-type annotations provided by the original study were directly adopted, with both modalities sharing six annotated cell types.

Gene symbols from the scRNA-seq data were mapped to their corresponding protein markers using the STRING database[53]. During preprocessing, proteins with variance lower than 0.1 in the CODEX data and genes with variance lower than 0.5 in the scRNA-seq data were excluded. The intersection of the remaining mapped proteins was then used as model input, resulting in 32 shared proteins for the tonsil dataset.

For model training, only the scRNA-seq data were used. The trained model was subsequently applied to the CODEX data for inference, and prediction performance was evaluated using the corresponding CODEX protein measurements.

### Spatium tokenization, architecture and pretraining

#### Spatium tokenization

Spatial proteomics pretraining in Spatium is conducted on a unified dataset comprising 51 million cells across 70 human spatial proteomics studies, covering tumor, immune, and stromal compartments. We reconciled heterogeneous protein panels into a unified feature vocabulary of 239 markers across studies.

Spatial proteomics datasets differ substantially in protein panel composition and annotation across studies. Due to differences in experimental design, identical proteins may appear under different names (e.g., ACTA2, αSMA, SMActin), and the overlap of measured proteins between datasets is limited. As a result, protein features are not naturally aligned across studies and cannot be directly compared without preprocessing. In addition, spatial proteomics panels are not designed to measure the full proteome. Instead, they consist of curated marker sets selected to distinguish predefined cell types. This leads to a feature space that is sparse and weak, where only a subset of proteins combination carries strong signal for cell-type discrimination, while many others show limited variation in each dataset. These characteristics introduce differences in measurement scale and background signal across studies. As a result, raw expression values are influenced by study-specific factors and cannot be directly compared between datasets. This motivates the use of representations that depend less on absolute intensity values.

To address these challenges, we propose a rank-based tokenization strategy. Given a dataset D*_d_*, we first standardize protein names using a manually constructed alias mapping that defines a unified protein vocabulary V of |V| = 234, which is provided in the (Supplementary Table 2). Each dataset contains a subset of observed proteins V*_d_* ⊆ V, reflecting dataset-specific panel designs. We then perform dataset-level z-score normalization to mitigate inter-study intensity scale shifts, followed by cell-wise ranking of protein expression. For each cell *i*, let *x_i_* ∈ ℝ^|V|^ denote the normalized protein expression vector, where missing proteins are masked. We define a ranking operator *π*(·):

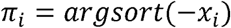

which produces an ordered index sequence such that:

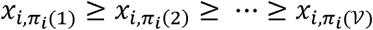

We then define a vocabulary mapping function *g*(·) that maps protein indices to unified protein identifiers:

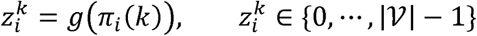

And convert them into token ids via:

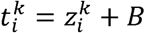

The final token sequence is constructed as:

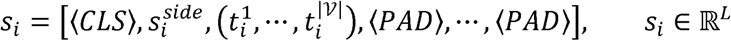

Where *B* denotes the protein token offset, where:

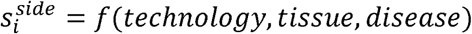

maps categorical metadata into discrete side-information tokens. All sequences are padded to a fixed length of *L* = 239 tokens to ensure uniformity across cells.

#### Spatium architecture and pretraining

Spatium adopts a Transformer-based encoder architecture for modeling ranked protein token sequences. The model consists of a 12-layer Transformer encoder, where each layer contains 16 self-attention heads. The hidden dimension is set to *d* = 256, and the feed-forward network dimension is 512. We denote the number of model parameters as a function of these hyperparameters, with a fixed vocabulary size of 278 vocabularies and a maximum sequence length of 239 protein positions plus auxiliary tokens. Each input sequence *s_i_* ∈ *R^L^* is mapped to a continuous representation via a learnable embedding matrix:

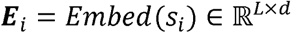

To model positional dependencies within ranked protein sequences, we introduce learnable positional embeddings:

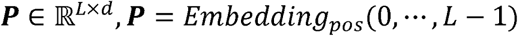

The final input representation is given by:

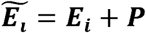

Unlike fixed sinusoidal positional encodings, we adopt learnable positional embeddings to allow the model to adaptively encode positional structure within protein token sequences. This design enables the Transformer to learn position-specific interaction patterns, particularly capturing how high-ranked proteins jointly define cellular identity within each sample.

Spatium is pretrained using a masked language modeling (MLM) objective over ranked protein token sequences[25]. Following a BERT-style corruption strategy, we randomly select 15% of non-special tokens (excluding CLS, PAD, and side-information tokens) for prediction. Among the selected tokens, 80% are replaced with a dedicated [MASK] token, 10% are substituted with random protein tokens, and the remaining 10% are kept unchanged. This design encourages the model to jointly learn contextual reconstruction and robustness to local perturbations in protein ranking sequences. Let s*_i_* denote the input token sequence for cell *i* and *s̃_ι_* is sequence with mask. The model is trained to reconstruct original tokens at masked positions. The training objective is defined as:

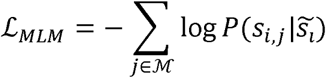

Where ℳ denotes the set of masked positions.

We optimize Spatium using the AdamW optimizer with weight decay regularization. The learning rate follows a stepwise schedule consisting of a linear warmup phase followed by cosine decay. Spatium is trained on two NVIDIA Tesla V100 24GB GPUs for approximately 16 hours. To improve I/O efficiency, all tokenized sequences are stored in Zarr format, enabling fast random access during training. Mixed-precision training FP16 is employed to reduce memory consumption and accelerate throughput. Model parameters are initialized using Xavier initialization for linear layers, while biases are initialized to zero. The training procedure operates at the step level, with the learning rate scheduler updated after each optimization step.

#### Cell representation

Each cell is represented as a ranked protein token sequence augmented with side-information tokens. Like sentence representation in natural language models, the contextual representation of a cell is obtained by aggregating token-level representations produced by the Transformer encoder. Although pooling operations such as mean pooling can be applied, we adopt a dedicated 〈*CLS*〉 token as the global representation of each cell. The 〈*CLS*〉 token is prepended to every input sequence, allowing the Transformer to learn an adaptive aggregation of contextual information through self-attention. The hidden state corresponding to the 〈*CLS*〉 position in the final encoder layer is extracted as the cell representation

For downstream tasks requiring a global characterization of cellular identity, including cell type annotation, spatial niche identification, and multi-modal inference, the 〈*CLS*〉 representation is directly used as the cell embedding for subsequent prediction heads. For protein imputation, however, the objective is to reconstruct the expression of individual proteins rather than produce a single cell-level embedding. Therefore, instead of using the 〈*CLS*〉 token, we utilize the contextualized representations of the ranked protein tokens. Specifically, the hidden representations corresponding to protein token positions are projected independently to predict protein expression values:

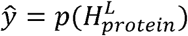

where 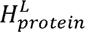 denotes the final-layer hidden representations of all protein tokens, and *p*(·) is a shared linear projection. This design preserves protein-level contextual information and enables the model to infer the relative expression of each protein, making it more suitable for panel expansion than a single pooled cell representation.

### Spatium downstream tasks

#### Cell type annotation and few-shot learning

We evaluate Spatium on the task of spatial cell type annotation under a few-shot learning setting. The pretrained Spatium model is used to initialize all model parameters, while the classification head is randomly initialized and trained from scratch. During fine-tuning, all parameters of the pretrained Transformer are updated unless otherwise specified. The model is optimized using a standard cross-entropy loss over annotated training cells. To evaluate model performance under limited annotation scenarios, we construct few-shot learning tasks by randomly sampling a small number of labeled cells per cell type from the reference set. Specifically, we vary the number of labeled samples per class *k* ∈ {1,2,5,10,20,50,100} to systematically evaluate performance under different annotation budgets.

For each setting, the classifier head is trained on the sampled support set and evaluated on a held-out query set consisting of spatially distinct regions. Performance is reported using accuracy and macro-F1 score, averaged over multiple random sampling runs to reduce variance.

#### Spatial niche analysis

To characterize higher-order spatial organization of tissues, we formulate a spatial niche composition prediction task, where the goal is to model the cell-type composition of local cellular neighborhoods. For each cell, we construct a spatial neighborhood by identifying its *k*-nearest neighboring cells within the same tissue section. We then compute a cell-type composition vector over the neighborhood, where each dimension represents the proportion of a specific cell type. Formally, for cell *i*, its neighborhood composition is defined as:

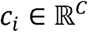

where *C* is the number of cell types. Each entry 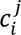 corresponds to the normalized frequency of cell type *j*within the neighborhood of cell *i*. Cell types do not present in the neighborhood are assigned to zero values.

We train Spatium to predict neighborhood composition vectors from learned cell representations. A regression head with randomly initialized parameters is applied to the cell embedding to output the predicted composition. The model is optimized using a mean squared error (MSE) loss. This objective encourages the model to capture spatially contextualized cellular interactions encoded in local neighborhood structures. After training, we extract the learned cell embeddings and construct a k-nearest neighbor graph based on cosine similarity in the embedding space. We then apply the Leiden algorithm[54] to identify spatially coherent clusters, which we interpret as latent spatial niches.

### Protein Intensity Imputation

To evaluate the capability of Spatium in recovering quantitative protein abundance, we formulate protein imputation as a continuous regression task. Unlike cell-level prediction tasks, protein imputation requires reconstructing the abundance of individual proteins while eserving both their relative ordering and protein-specific expression characteristics.

In addition to the ranked protein token sequence, we provide two auxiliary supervision signals derived from the original protein expression matrix. First, continuous protein intensities are quantile normalized within each tissue section to reduce technical variation in measurement scales while preserving relative abundance relationships. Let *X* ∈ ℝ*^N^*^×V^ denote the original protein expression matrix with quantile normalization, where *N* is the number of cells and V is the number of proteins and *X* will serve as the regression target during fine-tuneing. Second, to capture protein-specific abundance distributions, we discretize the normalized expression of each protein independently into *K* = 50 quantile bins across all cells within each tissue section. Specifically, for protein *p*, each cell is assigned a bin label *b_ip_* ∈ {0, ⋯, *K* − 1} forming a protein-wise bin matrix. This auxiliary representation captures the empirical abundance distribution of each protein and provides distribution-level supervision during optimization. Unlike cell-level downstream tasks that utilize the 〈*CLS*〉 representation, protein imputation directly operates on contextualized protein token representations. The contextualized representations corresponding to the ranked protein token sequence are projected independently through a shared linear layer to predict the continuous abundance associated with each ranked protein token:

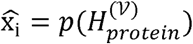

Where *x̂_ι_* ∈ ℝ*^P^* denotes the predicted protein abundance vector for cell *i* and *p*(·) denotes the shared linear layer.

To accurately reconstruct protein abundance, we optimize three complementary objectives. First, an L1 reconstruction loss preserves absolute protein intensities:

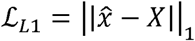

Second, since downstream biological interpretation of spatial proteomics primarily relies on the relative abundance of marker proteins rather than absolute fluorescence values, we introduce a ranking loss to preserve the ordering of protein expression within each cell:

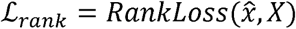

where the rank loss penalizes mismatches in pairwise ordering between predicted and ground-truth protein expressions.

Finally, protein abundances exhibit markedly different global expression distributions. While some proteins are broadly expressed across most cells, others are highly specific and remain nearly silent outside cellular populations. To encourage predicted abundances to remain consistent with these protein-specific distributions, predicted continuous values are first transformed into soft probability distributions over quantile bins using Gaussian kernel smoothing and then optimized against the ground-truth bin assignments using a Kullback–Leibler (KL) divergence loss:

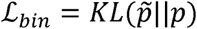

where *p̃* denotes the predicted soft bin distribution and *p* is the one-hot encoded ground-truth quantile bin.

The final optimization objective is defined as:

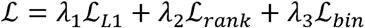

This multi-objective formulation jointly constrains continuous reconstruction accuracy, preservation of relative protein abundance, and consistency with protein-specific expression distributions, enabling Spatium to recover biologically meaningful protein intensity profiles from contextualized protein representations.

#### Multi-modal inference

For cross-modality cell type prediction, we adopted the same fine-tuning protocol as the cell type annotation task. Specifically, the pretrained Spatium backbone was initialized using pretrained weights, while the classification head was randomly initialized.

To enable compatibility between scRNA-seq and spatial proteomics, gene symbols from the scRNA-seq data were first mapped to their corresponding protein names using the unified protein vocabulary described in Section Human tonsil and pancreas dataset, after which only proteins shared by both modalities were retained. During fine-tuning, only the scRNA-seq reference cells with annotated cell types were used for supervised training. The trained classifier was subsequently applied to the paired CODEX dataset for cell type prediction without further modification.

To further explore the integration of histopathology and spatial proteomics, paired H&E whole-slide images were incorporated during fine-tuning. The H&E images were first spatially registered to the corresponding CODEX images. Image features were then extracted using the pretrained Prov-GigaPath[49] model. Each image was partitioned into overlapping patches, and patch-level visual embeddings were obtained using the pretrained vision encoder. Cell-level image representations were subsequently generated by bilinearly interpolating the embeddings of all overlapping patches covering each cell location.

During fine-tuning, the image embeddings were projected into the same latent space as the Spatium cell representations through a learnable projection layer. Cross-modal guidance was obtained by computing attention weights between protein-derived cell embeddings and image embeddings, producing image-guided contextual representations. The guided representations were fused with the original Spatium embeddings and optimized using a feature reconstruction objective, encouraging consistency between protein-derived and histology-derived representations while preserving the pretrained cellular representation.

#### Spatium simulation data construction

We construct a synthetic dataset by interpolating between two cell populations selected from CODEX data, corresponding to NK cells and CD66^+^ enterocyte cells, denoted as *C_NK_* and *C_CD_*_66_. These two populations are selected based on observed classification confusion matrix in the human intestine few-shot cell type annotation.

For each population, we compute a representative token sequence by taking median position over tokenized cell sequences within the corresponding class. Let *r_NK_* and *r_CD_*_66_ denote the resulting median token sequences. We construct a mixed rank template via linear interpolation:

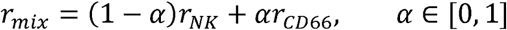

The resulting *r_mix_* is treated as a perturbation template. For each cell sequence *s_i_* in the dataset, we apply the perturbation by combining its original token sequence with *r_mix_* to obtain a modified sequence *sȃ_ι_*. The same template is applied uniformly across all cells. For each *α*, this procedure yields a perturbed dataset by applying the same mixed template to all cell token sequences. As *α* increases, the resulting token sequences progressively shift their marker composition toward the CD66⁺ enterocyte population in the token space.

### Spatium benchmarking

#### Few-shot cell type annotations

We evaluate Spatium and competing methods under a few-shot cell type annotation setting. For each dataset, we construct a reference set by uniformly sampling *k* = {1,2,5,10,20,50,100} cells per cell type from the annotated training split. The same reference and query splits are used across all methods to ensure fair comparison.

For Spatium fine-tuning, model parameters are initialized from the pretrained checkpoint. A task-specific classification head is randomly initialized and trained jointly with the encoder. All protein tokens are used as input. The learning rates are set to 5 × 10^−4^ for the encoder and 1 × 10^−3^ for the classification head.

For Spatium linear probing, the pretrained encoder is frozen and used to extract cell embeddings. A linear regression classifier is trained on reference embeddings and evaluated on query embeddings.

For Nicheformer[28], we use the pretrained weights from the official implementation via Hugging Face. Protein panels are mapped to gene symbols following the provided preprocessing pipeline, resulting in 44 shared features. Default hyperparameters are used, including a maximum sequence length of 1500. Cell embeddings are extracted from the final hidden layer and used as input to a linear regression classifier for evaluation.

For the PCA baseline, principal components are computed using all protein features. Since the number of reference samples in the *k* = 1 setting is 25, we fix the number of principal components to 25 to satisfy the rank constraint of PCA. PCA is independently fitted on the reference set and applied to both reference and query sets. The resulting embeddings are used to train a linear regression classifier.

For CytoVI[30], we follow the official preprocessing pipeline, including arcsine transformation and scaling of protein expression values. The model is trained on the reference set using default training settings, including the default warmup schedule and early stopping criterion. For label transfer, we use the provided impute_categories_from_reference function for inference on the query set.

#### Spatial niche analysis

For spatial niche analysis, we evaluate all methods under a unified regression framework. The task is formulated as predicting neighborhood-level cell type composition vectors.

For Spatium, the pretrained encoder is initialized from the pretrained checkpoint and fine-tuned with a randomly initialized regression head. The learning rates are set to 1 × 10^−4^for the encoder and 1 × 10^−3^for the regression head.

For CytoVI, we use the embeddings generated with the official implementation and default configuration. A regression head with identical architecture and optimization settings as Spatium is trained on top of the frozen embeddings.

For PCA, embeddings are generated using the first 30 principal components. A regression head with the same architecture and training configuration as Spatium is applied on top of the PCA embeddings.

Across all methods, the regression head is trained to predict the neighborhood cell type composition vector using a mean squared error (MSE) loss, ensuring a consistent evaluation protocol.

## Data availability

The datasets used for model fine-tuning and downstream analyses will be made publicly available upon publication of this work. Processed data and the corresponding metadata will be deposited in a public repository and Spatium github documentation, with accession numbers and download links provided in the final published version of the manuscript.

## Code availability

The code for Spatium, including model implementation, pretraining weights, and all downstream documentations, is publicly available at https://github.com/ploughhh/Spatium,

## Acknowledgment

We would like to thank all our colleagues and friends who provided advice on this study. We would also like to thank the researchers who selflessly shared their data. This work was partially supported by the National Institutes of Health [R01LM014156, R01GM153822, R01CA241930, R01AA032723-01 to X.Z], the National Science Foundation [2217515, 2326879 to X.Z], the Cancer Prevention and Research Institute of Texas [RP250043 to X.Z], and the Dementia Prevention and Research Institute of Texas [P018230 to J.L]. Dr. Liyu Huang is supported by the National Natural Science Foundation of China under grant Numbers 82227802 and 62373292. The funders had no role in study design, data collection and analysis, decision to publish or preparation of the manuscript.

