## Extended Data Figure and Supplementary Files. for "Spatium: A Protein Language Foundation Model for Spatial Proteomics": Supplementary Material.docx

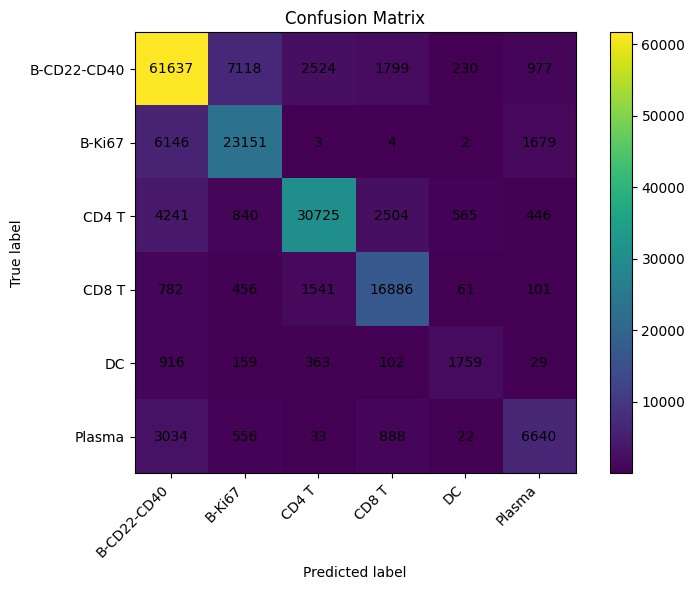

**Fig. 1 Confusion matrix of scRNA -Spatial Proteomics label transfer**.

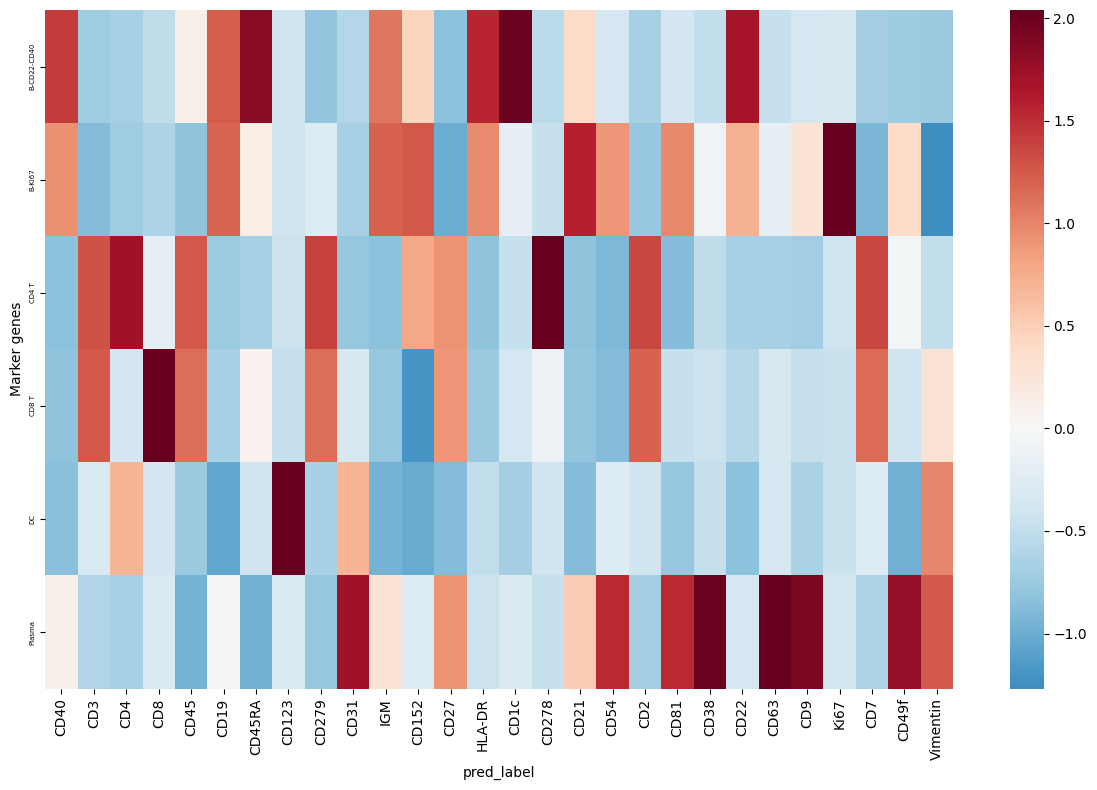

**Fig. 2 Marker distribution in all cell types.**
