## Supplementary figures and images for "Spatium: A Protein Language Foundation Model for Spatial Proteomics"

### Extended Data Fig 1.png

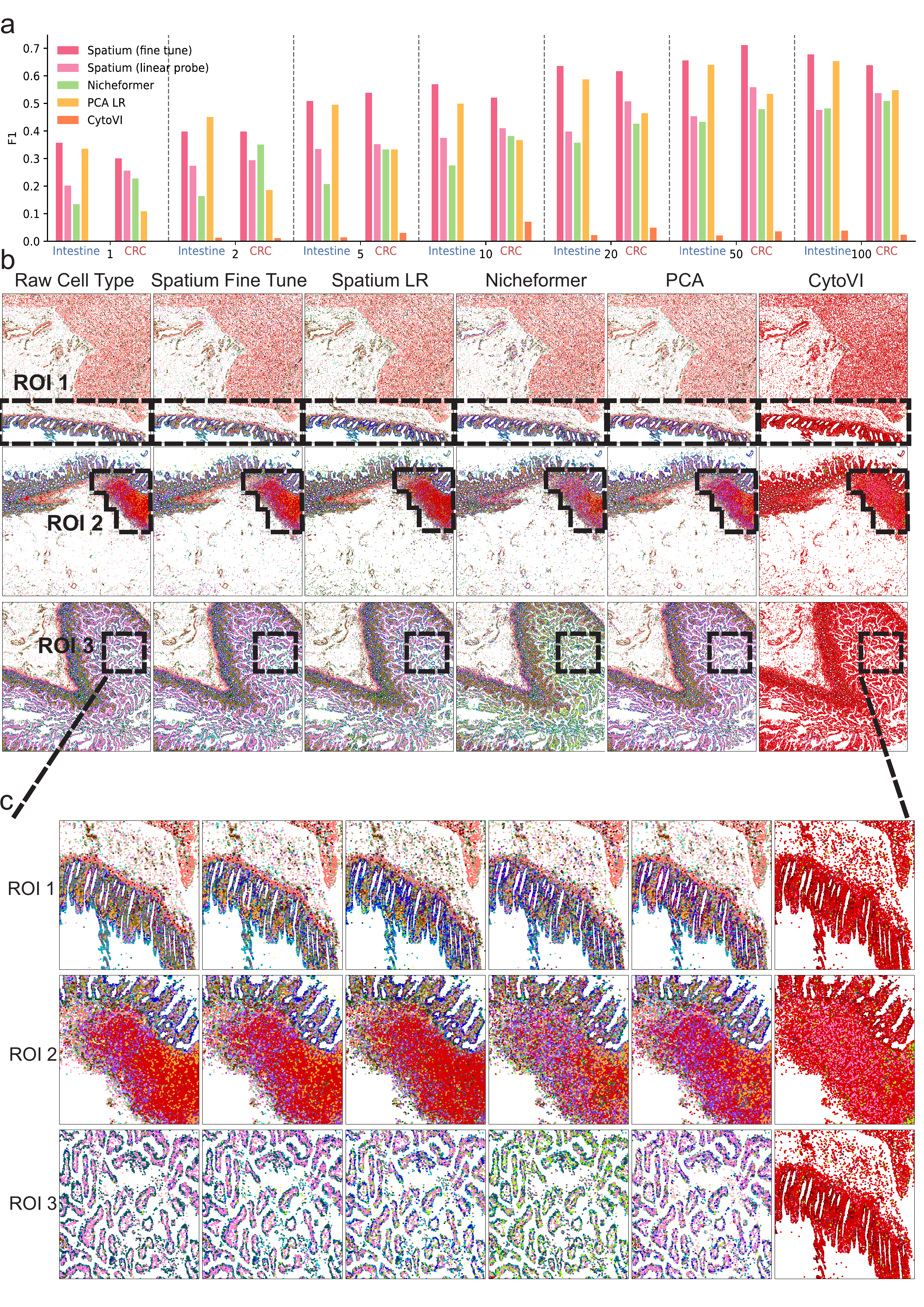

### Extended Data Fig 2.png

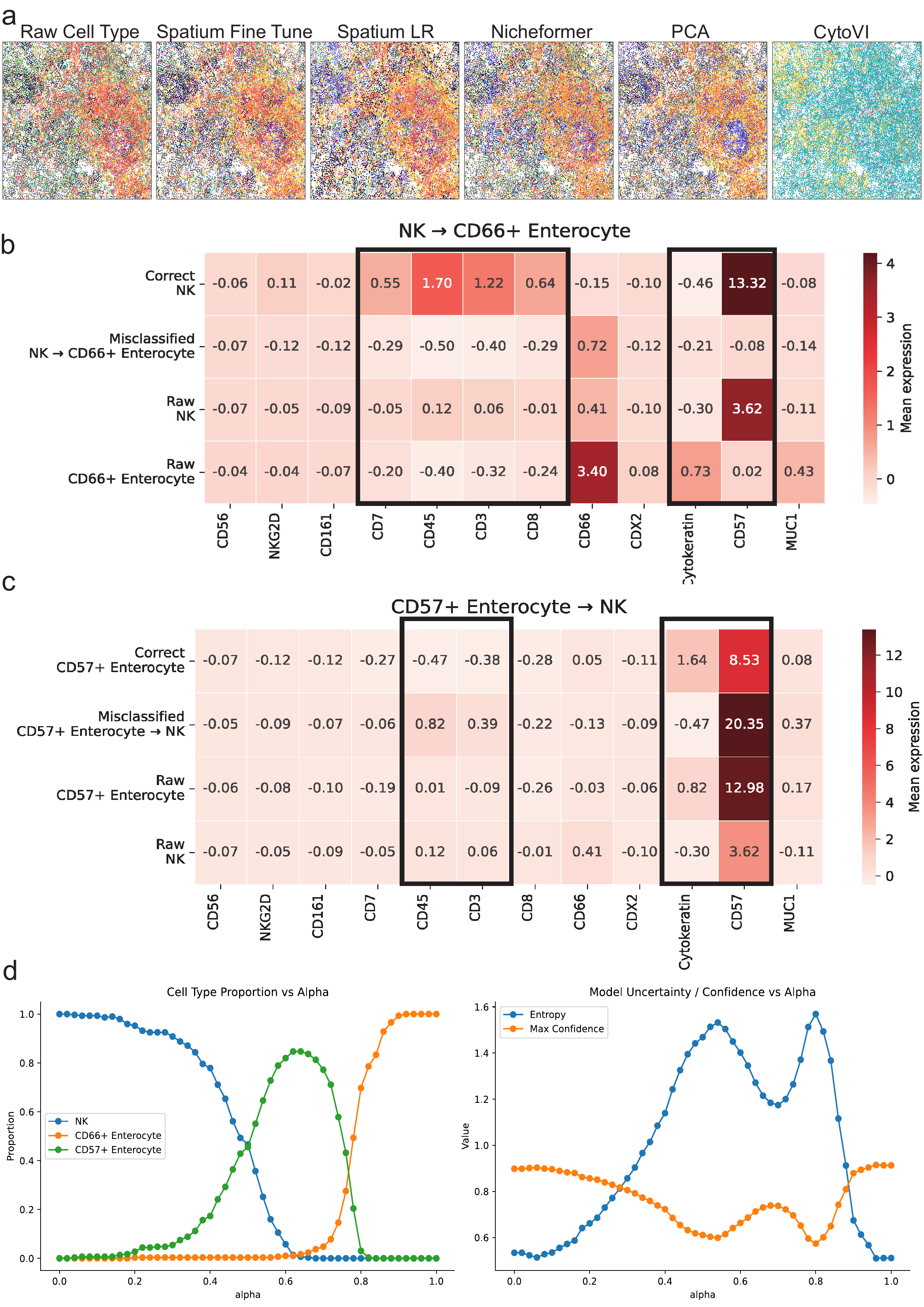

### Extended Data Fig 3.png

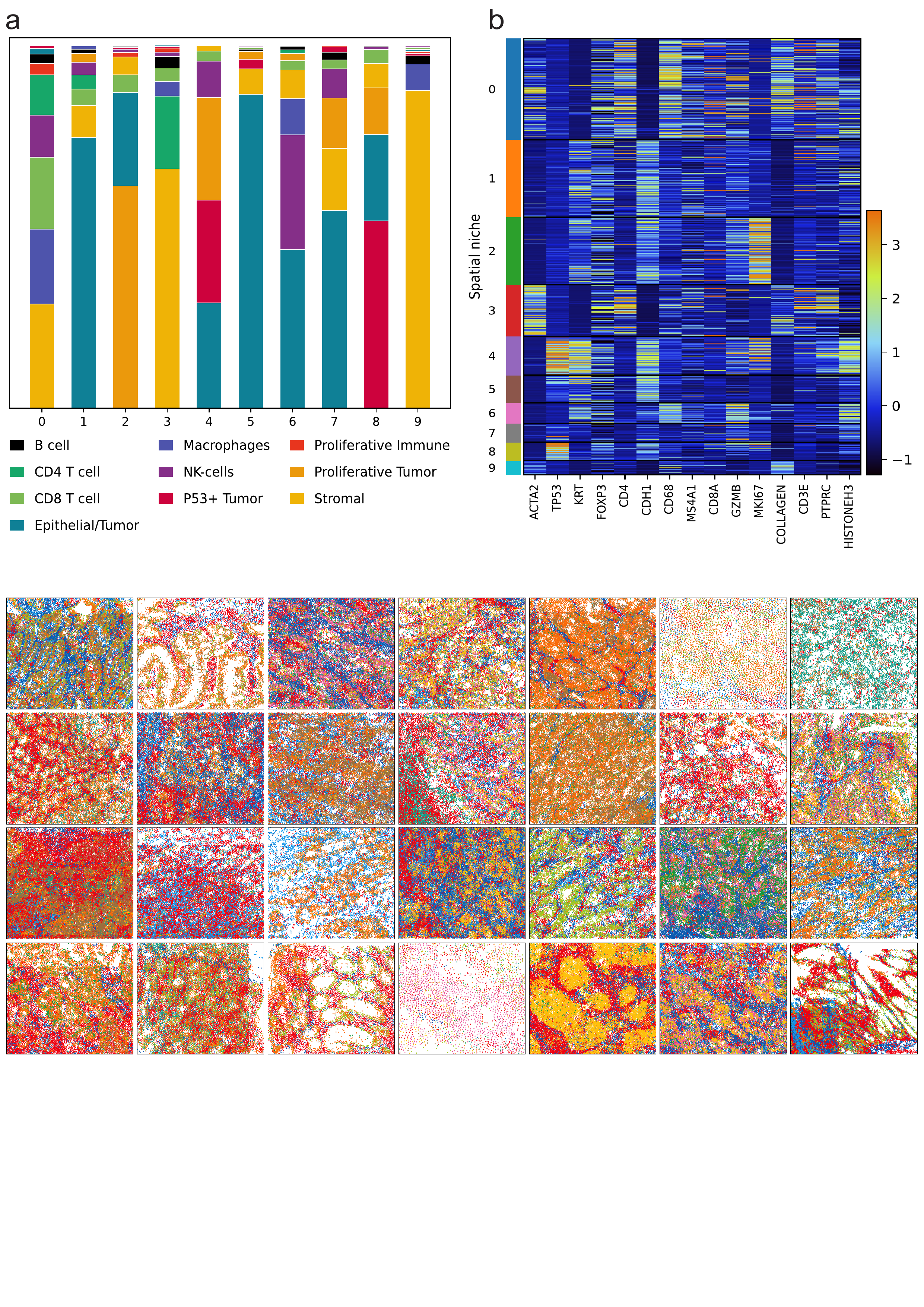

### Extended Data Fig 4.png

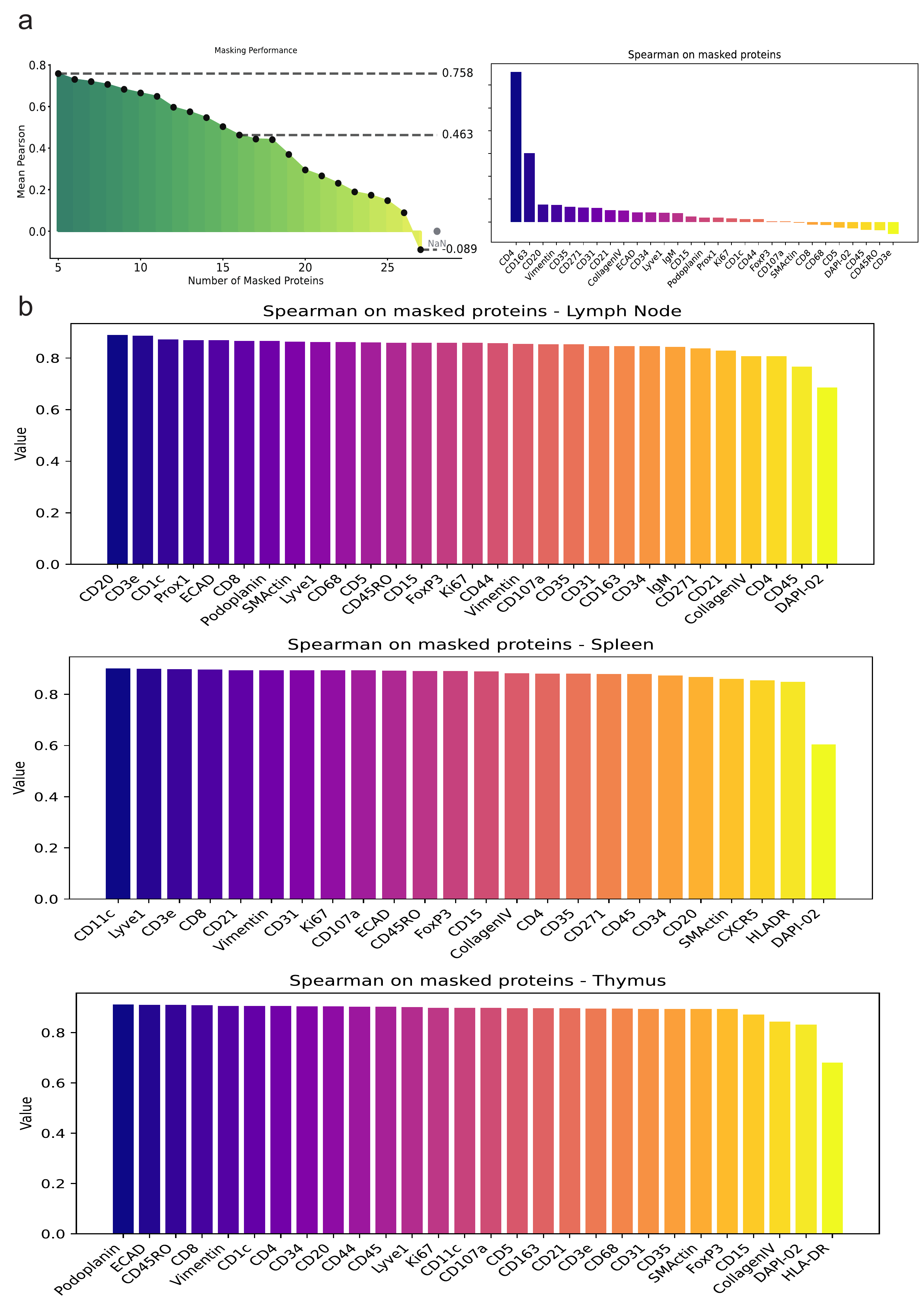

### Extended Data Fig 5.png

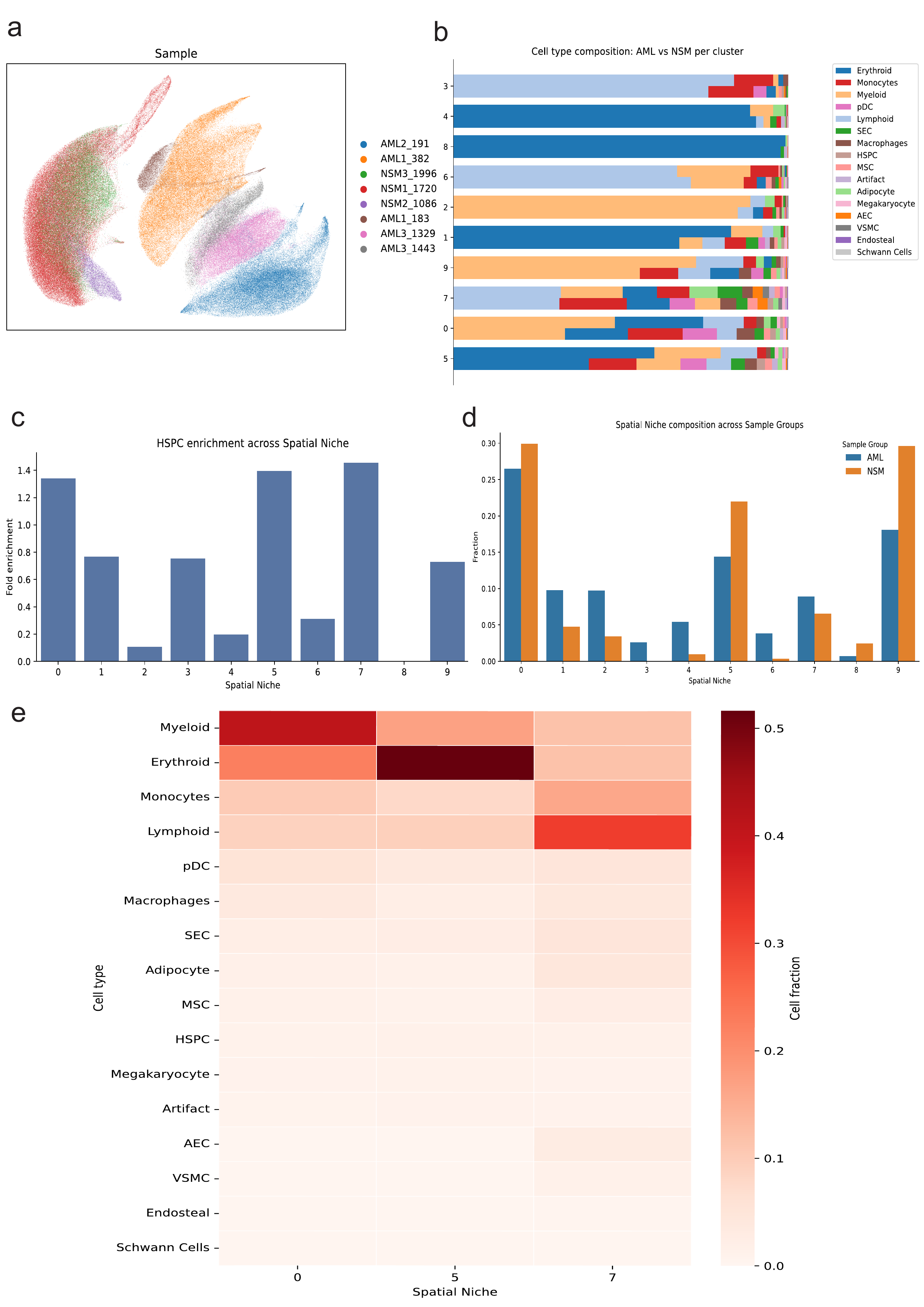
